# Yerba mate (*Ilex paraguariensis*) genome provides new insights into convergent evolution of caffeine biosynthesis

**DOI:** 10.1101/2023.09.08.556846

**Authors:** Federico A. Vignale, Andrea Hernandez Garcia, Carlos P. Modenutti, Ezequiel J. Sosa, Lucas A. Defelipe, Renato R.M. Oliveira, Gisele L. Nunes, Raúl M. Acevedo, German F. Burguener, Maximiliano Rossi, Pedro D. Zapata, Dardo A. Marti, Pedro A. Sansberro, Guilherme Oliveira, Madeline N. Smith, Nicole M. Dubs, Satish Nair, Todd J. Barkman, Adrian G. Turjanski

## Abstract

Yerba mate (*Ilex paraguariensis*) is an economically important crop marketed for the elaboration of mate, the third-most widely consumed caffeine-containing infusion worldwide. Here we report the first genome assembly of this species, which has a total length of 1.06 Gb and contains 53,390 protein-coding genes. Comparative analyses revealed that the large yerba mate genome size is partly due to a whole-genome duplication (Ip-α) during the early evolutionary history of *Ilex*, in addition to the hexaploidization event (γ) shared by core eudicots. Characterization of the genome allowed us to clone the genes encoding methyltransferase enzymes that catalyse multiple reactions required for caffeine production. To our surprise, this species has converged upon a different biochemical pathway compared to that of its relatives, coffee and tea. In order to gain insight into the structural basis for the convergent enzyme activities, we obtained a crystal structure for the terminal enzyme in the pathway that forms caffeine. The structure reveals that convergent solutions have evolved for substrate positioning because different amino acid residues facilitate a different substrate orientation such that efficient methylation occurs in the independently evolved enzymes in yerba mate and coffee. While our results show phylogenomic constraint limits the genes coopted for convergence of caffeine biosynthesis, the x-ray diffraction data suggests structural constraints are minimal for the convergent evolution of individual reactions.

## Introduction

In the genomic era, hundreds of plant species have had their nucleotide sequences determined to provide unprecedented insight into the genetic basis of many traits. One of the few species of economic importance for which no genomic data exists is *Ilex paraguariensis* var. *paraguariensis* A. St. Hilaire (Aquifoliaceae), colloquially known as yerba mate (YM), which is a caffeinated diploid tree-species (2*n*=2*x*=40) endemic to the subtropical rainforests of South America^1^. The dried leaves and twigs of this dioecious evergreen are used to prepare a traditional infusion named mate, or chimarrão, widely consumed around the world. Approximately 300,000 ha are cultivated with this tree crop, with Argentina responsible for 80% of world-wide production^2^. The mate infusion has been shown to have numerous beneficial effects in humans including as an antioxidant^3–5^, antidiabetic^6, 7^, as well as central nervous system stimulant^8^, among others. Several bioactive compounds have been identified in YM that might be responsible for its effects, including terpenes, flavonoids, phenolics and methylxanthines^2^. Although its stimulant properties are best known and mostly related to caffeine content, little is known about the genetic and biochemical mechanisms of how YM synthesizes this, or any, of its important metabolites. Despite the recent release of three other *Ilex* genome sequences^9–11^, none of the species produce caffeine, making the genetic basis for convergent evolution of this trait in YM unclear.

Convergent evolution has occurred throughout the tree of life and is particularly rampant in plants^12^ where examples of repeated origins of morphological^13^, anatomical^14^, physiological^15^, and biochemical^16^ traits have been documented. Caffeine (CF) is a xanthine alkaloid that has independently evolved no less than six times across angiosperms and has implications for pollination, insect defence and allelopathy^17, 18^. There are multiple biosynthetic routes to caffeine possible within the xanthine alkaloid network (Fig. 1). Within the Rosid genera *Theobroma*, *Paullinia*, and *Citrus*, sequential methylation of xanthine (X), 1- and/or 3- methylxanthine (1X, 3X) and, finally, either theophylline (TP) or theobromine (TB) leads to caffeine^19^. In contrast, the Asterids, *Coffea* and *Camellia*, appear to sequentially methylate xanthosine (XR), 7-methylxanthine (7X), and theobromine (TB) to yield caffeine^20, 21^ (Fig. 1). Regardless of which pathway is utilized, species differ in terms of which SABATH enzyme family members were convergently recruited to synthesize caffeine: Xanthine Methyltransferase (XMT) is used by *Citrus* and *Coffea* while the paralogous Caffeine Synthase (CS) is used by *Camellia*, *Theobroma* and *Paullinia*^19, 22, 23^ (Fig. 1). Convergence appears to also extend to the mutational level, since different amino acid replacements to homologous regions of CS and XMT enzymes appear to govern the evolution of substrate preference switches^24^. However, it remains unclear whether mutations lead to convergent three-dimensional protein structures to confer convergent substrate interactions and catalysis by the enzymes. Because XMT- and CS-type enzymes have been convergently recruited in both Rosids and Asterids to catalyse the same or different pathways, it suggests considerable evolutionary lability underlying caffeine production in plants^19^. As a result, it is difficult to predict what sets of genes, biochemical reactions, and structural properties might lead to the evolution of caffeine biosynthesis in YM, or any plant, *a priori*.

**Fig. 1.**
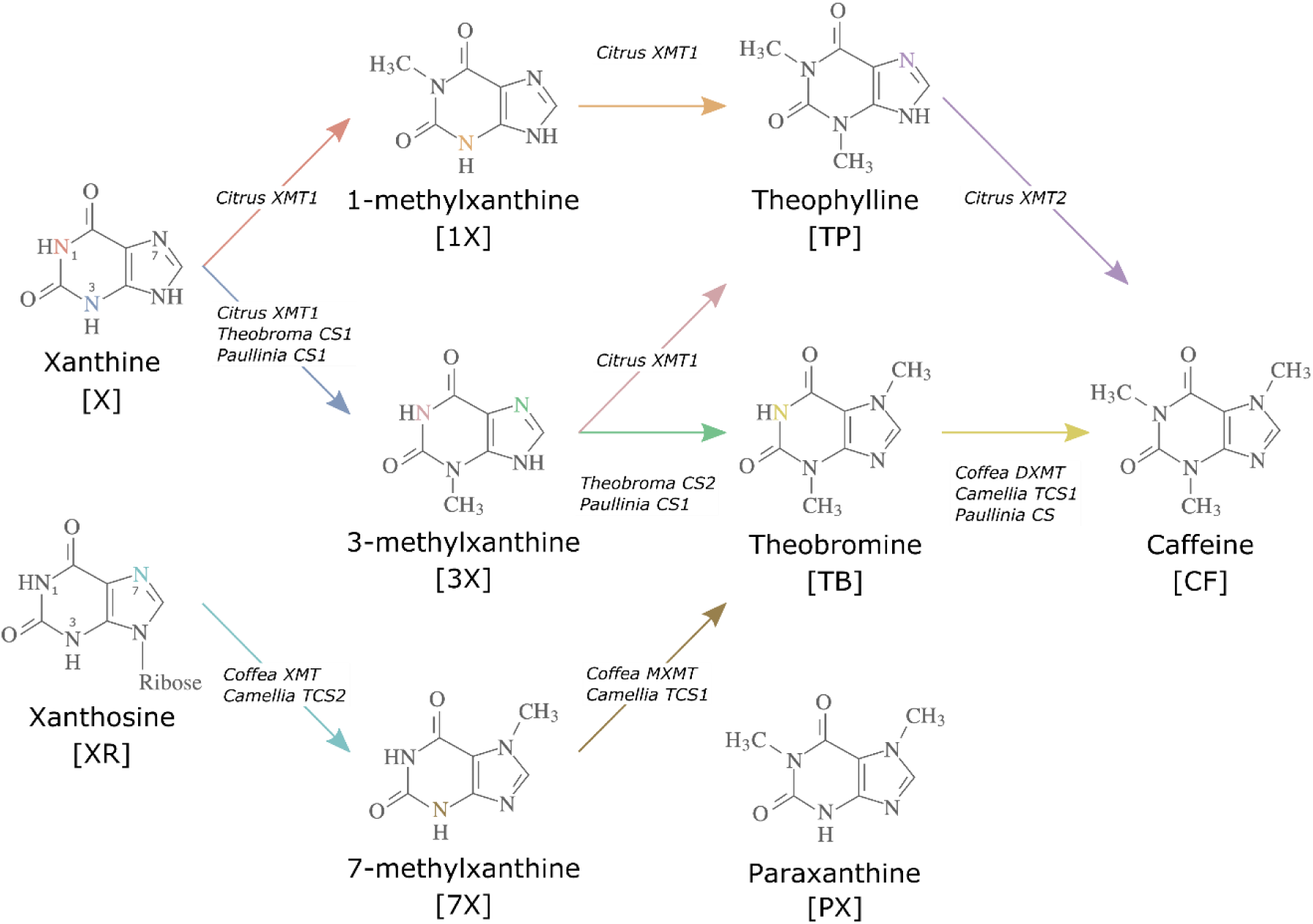
Biosynthetic routes to caffeine within the xanthine alkaloid network. CF, caffeine; PX, paraxanthine; TB, theobromine; TP, theophylline; 1X, 1-methylxanthine; 3X, 3- methylxanthine; 7X, 7-methylxanthine; XR, xanthosine; X, xanthine. Nitrogen atoms are coloured to match the arrows corresponding to the enzymes that methylate them. Adapted with permission from O’Donnell et al.^24^

Although some transcriptomic resources have been generated for YM^25–27^, a complete genome sequence has the potential to advance our understanding of the metabolic potential of this important crop and facilitate improvement. Here, we describe the first draft genome of YM and report on its composition, organization and evolution. The genomic sequence enabled us to uncover the genetic, biochemical and structural bases for convergent evolution of caffeine in YM. Our comparative analyses of caffeine-producing enzymes across angiosperms reveals how convergence may be the result of constrained evolutionary genomic potential but relatively unconstrained structural potential.

## Results and Discussion

### Yerba mate genome sequencing, assembly and annotation

The YM genome was sequenced combining Illumina and PacBio sequencing technologies. With Illumina sequencing, we generated ∼263.2 Gb of short reads from various DNA fragment sizes (350 bp, 550 bp, 3 kbp, 8 kbp and 12 kbp), while with PacBio sequencing, we generated ∼77.5 Gb of long reads. These reads represent ∼158.5- and ∼49.3-fold base-pair coverage of the genome, respectively (Supplementary Table 1). The total assembly length was ∼1.06 Gb and consisted of 10,611 scaffolds (≥1 kb) with an N50 length of ∼510.8 Kb (Supplementary Table 2). To assess the completeness of the assembly we aligned the available YM transcriptome reads^25–27^ and the YM genome short reads generated in this study with the genome assembly, establishing that 99.3% of the YM transcriptome reads and 99.5% of the YM genome short reads mapped to the assembly. This result suggests that we have assembled almost all available sequence data. The GC content of the genome assembly was 36.33% (Supplementary Table 2), similar to that of other eudicots (33.70-38.20 GC%)^28^ and almost identical to that of *Ilex polyneura* (36.08 GC%)^9^, *Ilex asprella* (36.25 GC%)^10^, and *Ilex latifolia* (36.44 GC%)^11^, the only three *Ilex* species with sequenced genomes. About 64.63% of the genome assembly was composed of repetitive sequences, of which ∼36.22% were retrotransposons, ∼1.80% were DNA transposons, ∼0.74% were simple repeats and ∼0.15% were low complexity regions. Long terminal-repeat (LTR) retrotransposons of the Gypsy and Copia families were the most abundant transposable elements, as observed in many sequenced plant genomes^29^, followed by long interspersed nuclear elements (LINEs) and hobo-Activator transposons, among others (Supplementary Table 3).

A total of 53,390 protein-coding genes were predicted in the genome, with a mean coding sequence length of 3,062 bp and 4.23 exons per gene. Of these, 41,483 (∼77.63%) could be annotated with GO terms, EC numbers or Pfam domains. In addition, we identified 4,530 non-coding RNA genes, including 2,670 small nucleolar RNAs, 815 transfer RNAs, 471 ribosomal RNAs, 348 small nuclear RNAs, and 226 micro RNAs (Supplementary Notes S1 and Supplementary Tables 4-6). To further assess the completeness of the assembly, we aligned the scaffolds with the KOG^30^ and DEG^31^ databases, determining that 98% of the core gene families from the KOG database and 97.5% of the *Arabidopsis thaliana* DEG subset were present in the assembly. Then, we performed a Benchmarking Universal Single-Copy Orthologs (BUSCO)^32^ assessment using the eudicot ODB10 database. Among 2,326 conserved single-copy genes, ∼96.20% were retrieved, of which ∼78.80% were complete and single copies, ∼17.40% were complete and in duplicates, ∼3.10% were fragmented and only ∼0.70% were missing. These results suggest that the coding region of the assembly is nearly complete. The number of estimated genes for YM is higher than the ca. 39,000 reported from the genome sequences of other *Ilex* species^9–11^. This could be at least partly due to the larger genome size of YM estimated from flow cytometry relative to the other species^33^.

### Evolutionary analysis of yerba mate genome provides evidence of whole-genome duplication in an early *Ilex* ancestor

Most plant lineages have experienced ancient polyploidization events followed by massive duplicate gene losses and genome rearrangements, which may have contributed to the evolution of developmental and metabolic complexity^34, 35^. Recent transcriptome-based analyses^36, 37^ reported an ancient polyploidization event in the *Ilex* lineage around 60 Ma (Cretaceous–Paleogene boundary), based on phylogenomic and synonymous substitution rate (K_s_) evidence. Evolutionary analyses of *Ilex polyneura*^9^ and *Ilex latifolia*^11^ genomes also provided evidence of a shared *Ilex*-specific whole genome duplication (WGD). As YM is the first American holly to have its genome sequenced, we performed synteny-based analyses of its genome to deepen our understanding of Aquifoliales evolution (Fig. 2 and Supplementary Fig. 1). The K_s_ distribution of YM paralogues (Fig. 2b) revealed a significant peak with a median K_s_ value of ∼0.37, not shared with the rest of the eudicot genomes analysed (Fig. 2b and Supplementary Fig. 1). This confirms the lineage-specific polyploidization event (Ip-α) previously reported in *Ilex*^9, 11, 36, 37^, in addition to the shared ancestral WGT-γ which is indicated by a median K_s_ value of ∼1.4 (Fig. 2b). A WGD in the common ancestor of *Ilex* species is further supported by 2:1 syntenic depth ratios between the YM genome and the coffee and grape genomes, which did not experience additional duplication events after the ancestral WGT-γ (Fig. 2c). In order to determine the age of Ip-α, we used two different phylogenies (Fig. 2a). The plastid genome phylogeny supports the monophyly of Aquifoliales as the first diverging clade of campanulids^38^; the alternative nuclear genome phylogeny supports *Ilex* in Aquifoliales I as an early branching lineage of lamiids^36^. With the former phylogeny, we estimated the age of the WGD event between 48.75-69.63 Ma while, with the latter, divergence was estimated at 49.43-70.62 Ma (Fig. 2a). Both estimates are consistent with that of Zhang et al. 2020 and validate the age of Ip-α near the origin of *Ilex*, which is estimated between 43-89 Ma^39^.

**Fig. 2.**
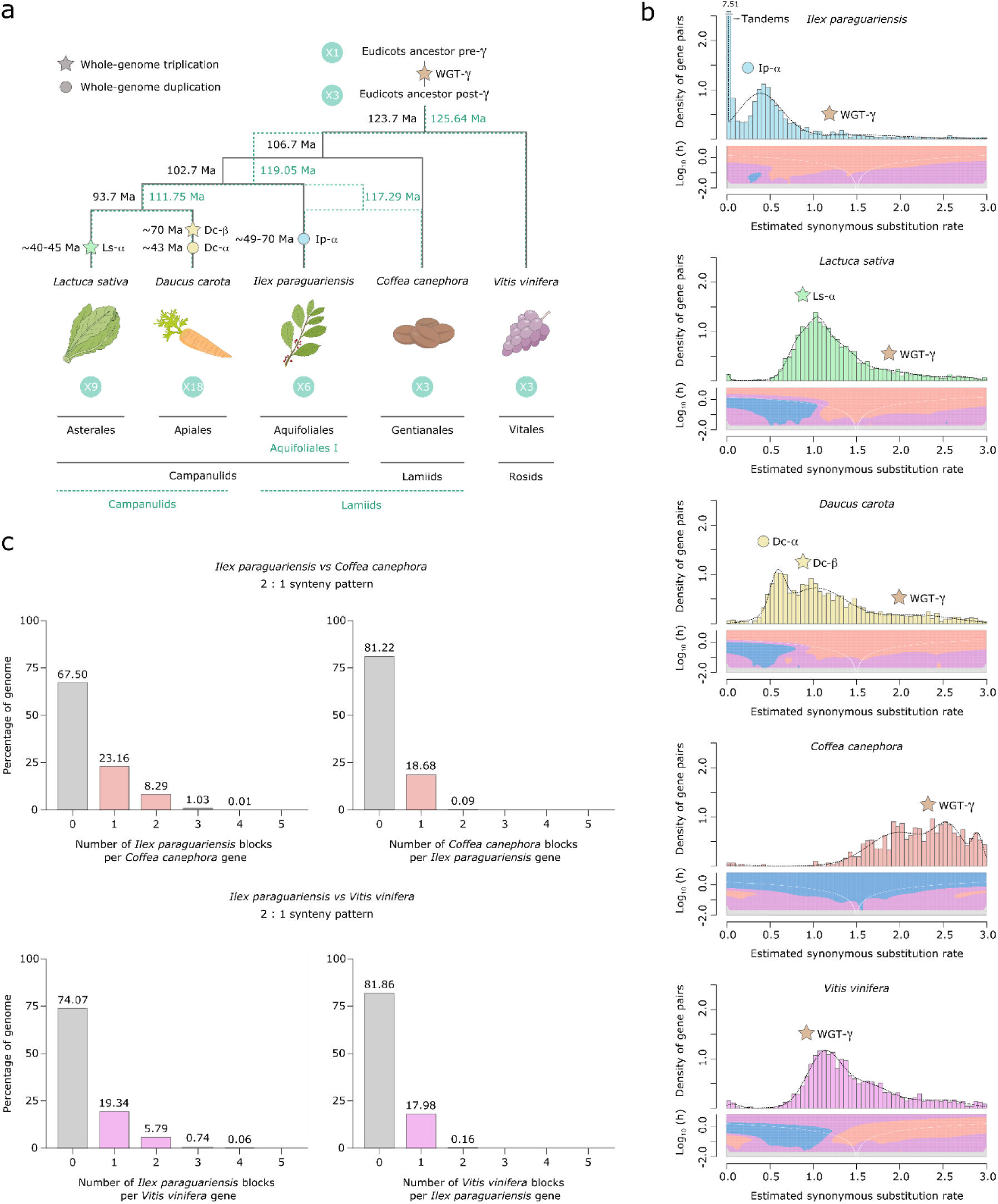
Yerba mate genome duplication history. **a**, Evolutionary scenario of the eudicot genomes of *Lactuca sativa*, *Daucus carota*, *Ilex paraguariensis*, *Coffea canephora* and *Vitis vinifera*, from their ancestor pre-γ. The plastid genome phylogeny is represented with solid black lines, while the multiple nuclear genome phylogeny is represented with green dashed lines. Paleopolyploidizations are shown with coloured dots (duplications) and stars (triplications). Divergence time estimates for the lineages, as well as age estimates for the *L. sativa* and *D. carota* paleopolyploidizations were obtained from the literature^36, 38, 100, 101^. Ma, million years ago. **b**, K_s_ distributions with Gaussian mixture model and SiZer analyses of *I. paraguariensis* (blue), *L. sativa* (green), *D. carota* (yellow), *C. canephora* (red) and *V. vinifera* (purple) paralogues. SiZer maps below histograms identify significant peaks at corresponding K_s_ values. Blue represents significant increases in slope, red indicates significant decreases, purple represents no significant slope change, and grey indicates not enough data for the test. **c**, Comparative genomic synteny analyses of *I. paraguariensis* with *C. canephora* and *V. vinifera*.

### Convergent evolution of caffeine biosynthesis in yerba mate

In order to determine the genes and biochemical pathway responsible for caffeine biosynthesis in YM, we used bioinformatic analyses to identify SABATH enzyme family members in the genome^19, 22, 23^. There appear to be 28 full-length SABATH genes in YM that encode members of the functionally diverse clades of the family, including SAMT^40^ and JMT^41^, among others (Fig. 3a). Our phylogenetic analysis showed that although the YM genome does not appear to encode XMT-type caffeine-producing enzymes like *Coffea* and *Citrus*, it does contain three recently and tandemly duplicated genes that encode CS-type enzymes, IpCS1- 3 (Fig. 3a, c). The duplicated IpCS1-3 are 86-91% identical at the amino acid level and are expressed at highest levels in caffeine-accumulating tissue (Fig. 3b)^19^. IpCS1-3 also appear to be of recent origin, since non-caffeine accumulating *Ilex* species only have a single gene or gene fragment in the syntenic region (Fig. 3c). In *Camellia*, *Theobroma* and *Paullinia*, recent duplications of the CS-type enzymes responsible for the successive steps of xanthine alkaloid methylation have also independently occurred (Fig. 3a)^19, 24^. Two other YM genes encode IpCS4 & 5, but these are not syntenic with IpCS1-3 and are not highly expressed in any tissues studied (Fig. 3b); therefore, we did not characterize them further.

**Fig. 3.**
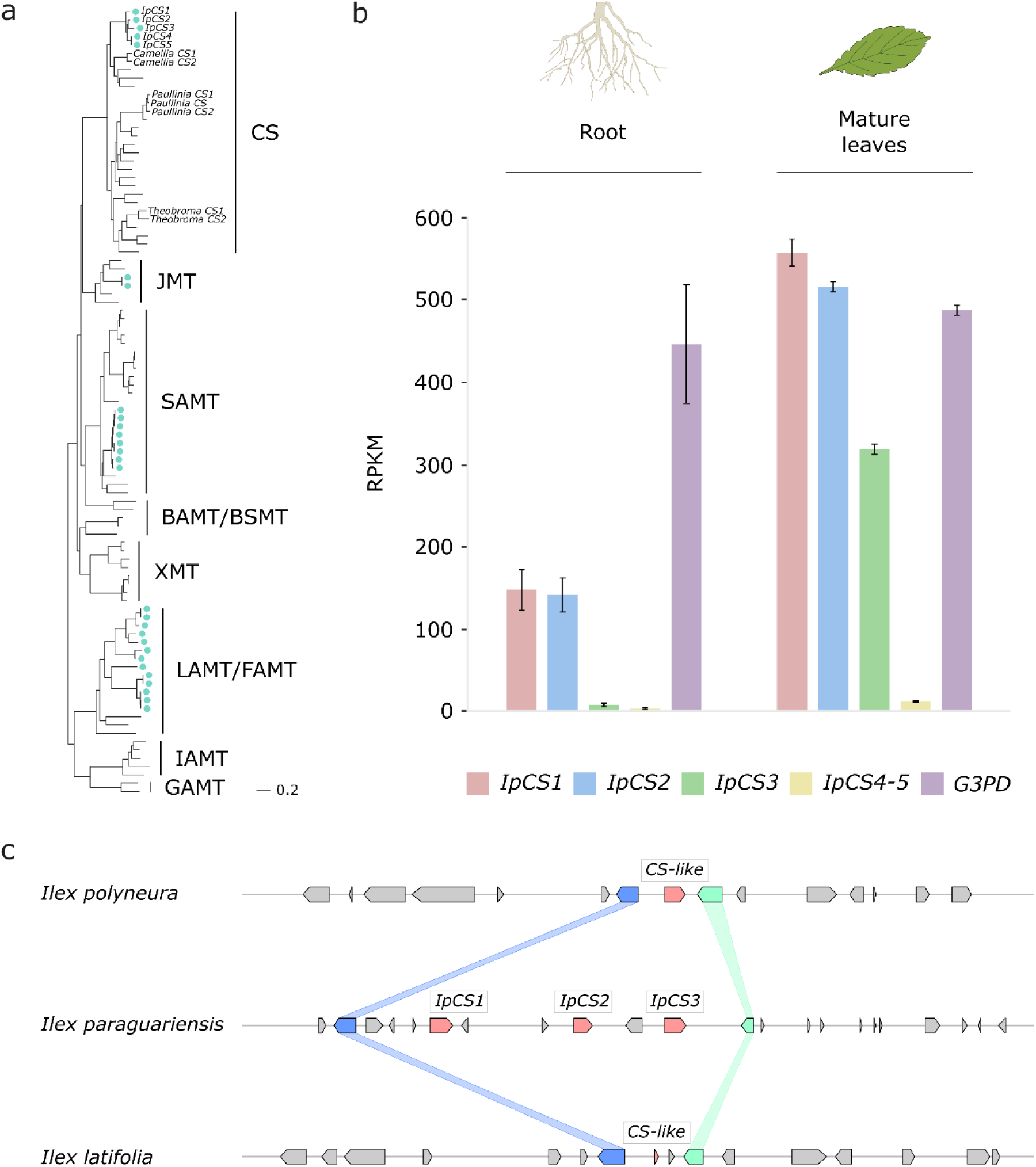
The yerba mate genome encodes three recently duplicated CS-type SABATH proteins that are expressed in caffeine-producing tissues. **a**, SABATH gene tree estimate (LnL = −34265.473) shows the placement of full length YM proteins (marked by blue-green dots) within clades that have published functions. GAMT, gibberellin MT; IAMT, indole-3-acetic acid MT; LAMT/FAMT, loganic/farnesoic acid MT; BAMT/BSMT, benzoic/salicylic acid MT; XMT, xanthine alkaloid MT used for caffeine biosynthesis in *Coffea* and *Citrus*; SAMT, salicylic acid MT; JMT, jasmonic acid MT; CS, caffeine synthase in *Theobroma*, *Camellia* and *Paullinia*. **b**, Gene expression analysis of IpCS1-5 as indicated by the relative abundance of YM transcriptome reads mapped to the IpCS1-5 transcripts. RPKM, reads per kilobase per million mapped reads. Error bars indicate standard deviation from the mean. Housekeeping gene: *G3PD*, Glyceraldehyde-3-phosphate dehydrogenase. **c**, Synteny-based analysis of the CS genomic region for *I. paraguariensis*, *I. polyneura* and *I. latifolia*.

To investigate the biochemical activities of the enzymes encoded by the three CS-type genes, we cloned them into bacterial expression vectors and determined heterologous protein functions. One enzyme, IpCS1, appears to primarily methylate X to catalyse the formation of 3X (Fig. 4). A second enzyme, IpCS2, shows activity only with 3X to produce TB, while a third enzyme, IpCS3, exhibits a preference to methylate TB to form CF (Fig. 4). Thus, collectively, these three enzymes appear capable of catalysing a complete pathway from xanthine to caffeine. The apparent *K_M_* for the preferred substrates of all three enzymes ranges from 85- 197 μM, and the *k_cat_* / *K_M_* estimates are comparable to those determined for other caffeine biosynthetic enzymes^24^ (Supplementary Table 7). Further evidence for this biosynthetic pathway has been reported by ^14^C xanthine tracer studies in young leaf segments of *I. paraguariensis* that showed radioactivity in 3X and TB in addition to CF^42^. A pathway from X→3X→TB→CF has also been reported for *Theobroma* and *Paullinia* using CS-type SABATH enzymes^19^. Like Huang et al. 2016, this represents another departure from the long-assumed pathway to caffeine biosynthesis (XR→7X→TB→CF) as reported in coffee and tea (Fig. 1). This instance in *Ilex* is particularly notable since YM is an Asterid, like coffee and tea. The fact that *Ilex*, *Theobroma* and *Paullinia* convergently recruited CS genes that independently duplicated and evolved to encode enzymes with similar substrate preferences to catalyse a common pathway to caffeine, in spite of their divergence more than 100 Ma^43^, is remarkable and suggests a high degree of genetic constraint governing the repeated origin of this trait.

**Fig. 4.**
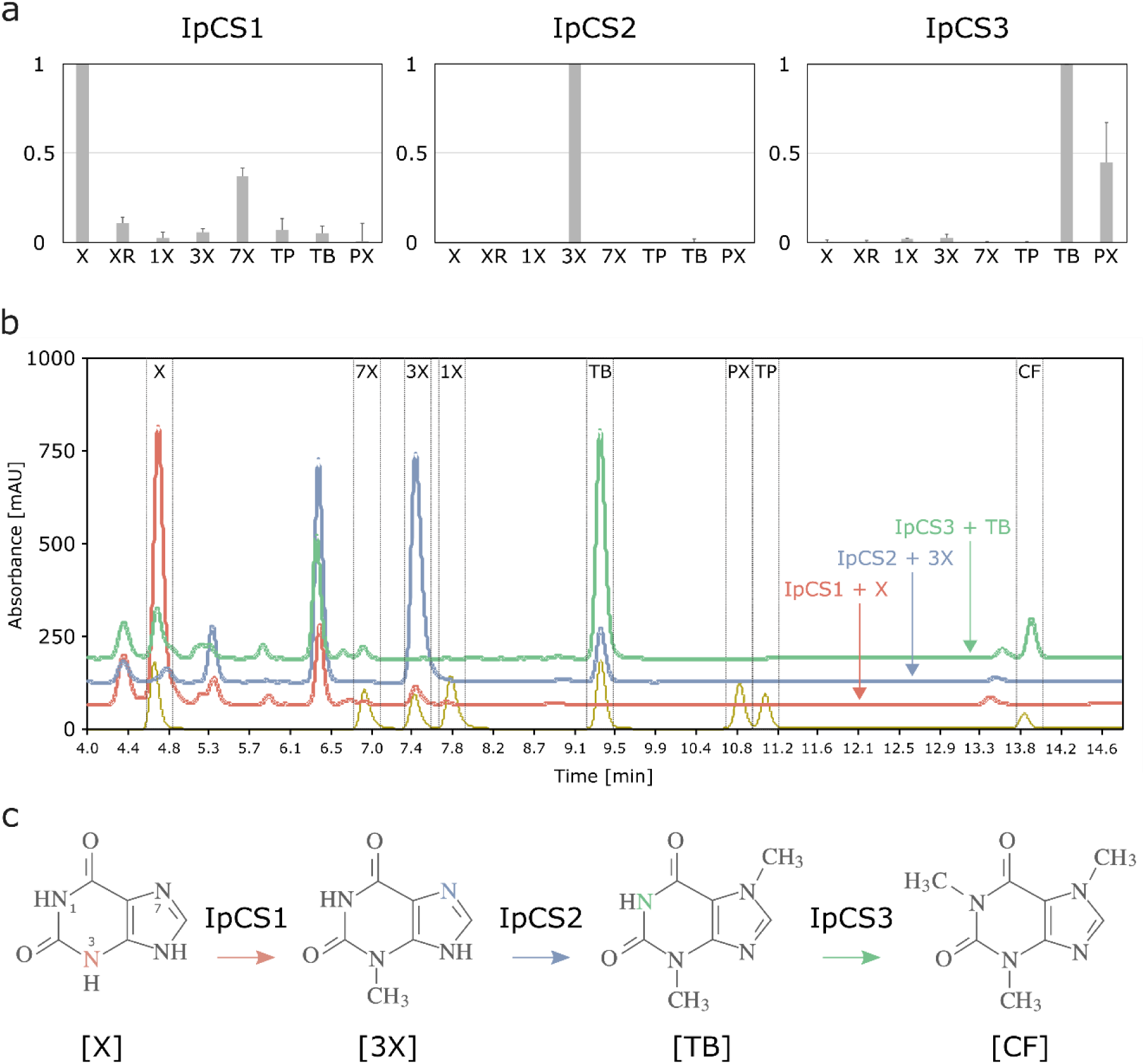
SABATH enzymes have evolved to catalyse the biosynthesis of caffeine in yerba mate. **a**, Relative enzyme activities of three CS-type SABATH enzymes with eight xanthine alkaloid substrates. **b**, HPLC traces showing products formed by three encoded CS-type enzymes. **c**, Proposed biosynthetic pathway for caffeine in yerba mate. X, xanthine; XR, xanthosine; 1X, 1-methylxanthine; 3X, 3-methylxanthine; 7X, 7-methylxanthine; TP, theophylline; TB, theobromine; PX, paraxanthine. Coloured atoms and arrows indicate atoms that act as methyl acceptors for a given reaction.

While the substrate preferences shown in Fig. 4 suggest pathway flux from X→3X→TB→CF, IpCS1 also shows secondary activity with 7X to produce TB and IpCS3 can catalyse the formation of CF from paraxanthine (PX) (Fig. 4a). Thus, flux through other branches of the xanthine alkaloid biosynthetic network (Fig. 1) cannot be excluded. However, it is not clear how 7X or PX would be produced *in planta* since none of the three enzymes studied here is capable of their formation; therefore, these secondary activities may not be physiologically relevant. In addition, it has been proposed that TP may also be a precursor to caffeine biosynthesis in *I. paraguariensis* based on radioisotopic feeding studies^42^, although its levels in plant tissues are 30-160 times lower than TB^44^. Our *in vitro* enzyme assays provide no experimental evidence for that biosynthetic route; however, it is possible that additional MT enzymes from the SABATH (or other) gene family not characterized in this study may perform such reactions. Alternatively, if the exogenously supplied TP was first catabolized to 3X in YM tissues, then the caffeine detected previously^42^ could have been synthesized via the route described above for IpCS2+IpCS3 (Fig. 4).

### The caffeine biosynthetic pathway in YM evolved from ancestral networks with different inferred flux

The fact that caffeine is produced within only one small lineage of *Ilex* that diverged and experienced CS gene duplication (Fig. 3) within the last 11 million years^39, 44^ indicates that the pathway has only recently evolved. The nature by which novel multistep biochemical pathways evolve is a central question in biology^45^. To investigate the caffeine pathway origin in YM, we used Ancestral Sequence Resurrection^46, 47^ to study AncIpCS1 & AncIpCS2, the ancestors of the three modern-day enzymes implicated in caffeine biosynthesis in YM (Fig. 5 and Supplementary Fig. 2-5). The ancestral enzyme, AncIpCS1, which gave rise to all three modern-day YM enzymes, exhibits highest relative activity with X, 3X and 7X (Fig. 5a). Methylation of 7X by AncIpCS1 occurred at the N3 position resulting in TB synthesis, whereas xanthine methylation occurred at either the N1 or N3 position to form 1X and 3X, respectively (Fig. 5b and Supplementary Fig. 6). AncIpCS1 was capable of methylation of 3X at N1 to produce TP, while methylation at the N7 position led to TB formation (Fig. 5b). These data demonstrate that, although AncIpCS1 could not produce caffeine, it could methylate xanthine alkaloids at 3 different positions of the planar heterocyclic ring structures and this combination of activities would have allowed for the ancestor of YM to produce a cocktail of 1X, 3X, TP and TB by flux through multiple branches of the xanthine alkaloid biosynthetic network with a single enzyme (Fig. 5b).

**Fig. 5.**
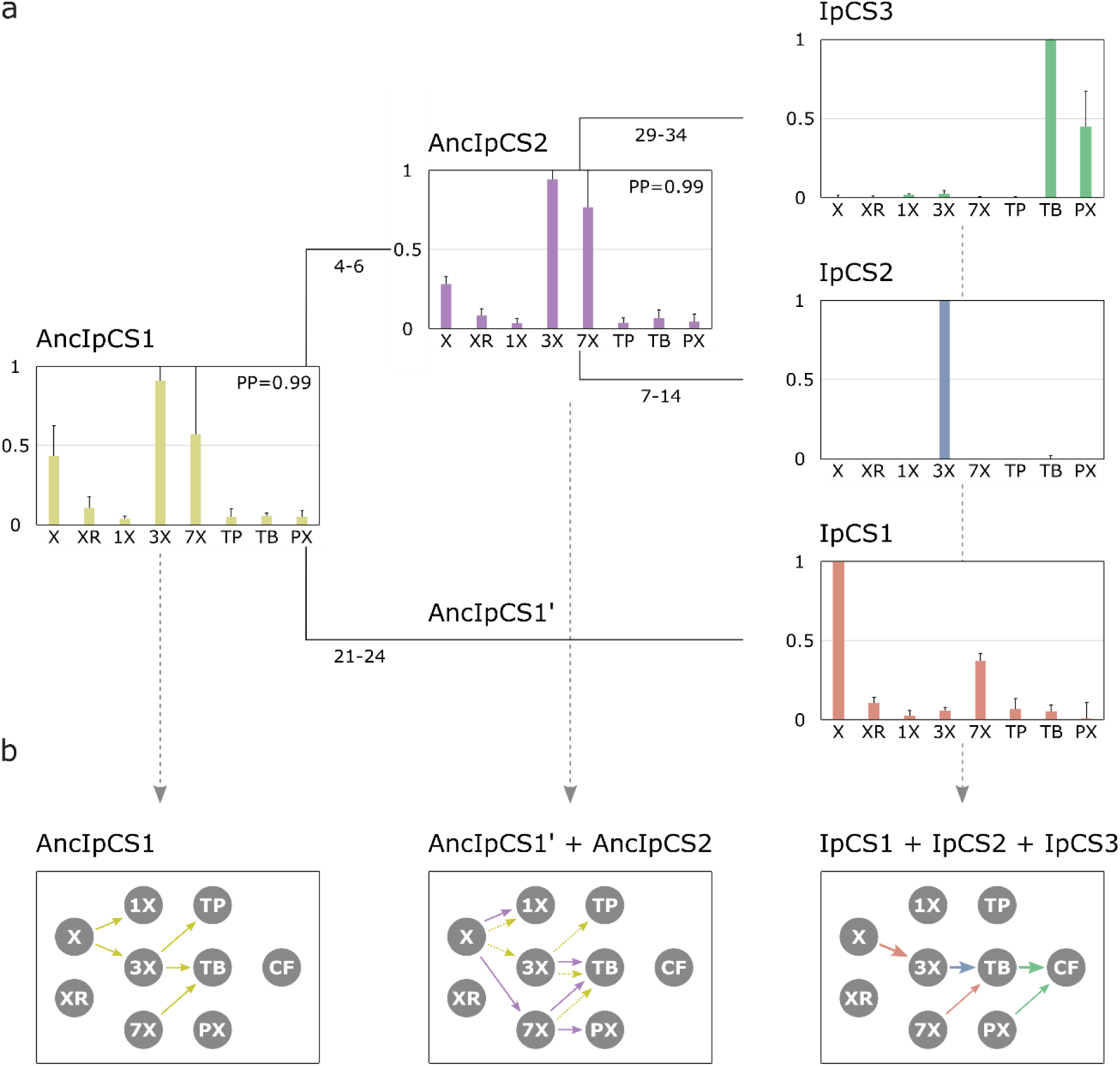
Ancestral sequence resurrection reveals ancestral xanthine alkaloid pathway flux. **a**, Simplified evolutionary history of three YM xanthine alkaloid-methylating enzymes and their two ancestors, AncIpCS1 and AncIPCS2. Average site-specific posterior probabilities for each ancestral enzyme estimate are provided. Numbers below each branch of the phylogeny represents the number of amino acid replacements between each enzyme shown. These two ancestral relative activity charts show the averaged activities of two allelic variants of each enzyme. **b**, Inferred pathway flux is shown for the antecedent pathways that could have been catalysed by the ancestral or modern-day combinations of enzymes that would have existed at three time points in the history of the enzyme lineage. Arrows linking metabolites are coloured according to the activities detected from each enzyme shown in panel a. Dotted arrows are shown for AncIpCS1’ because it is unknown what characteristics it would possess; it is assumed that it would have at least catalysed the formation of 3X from X since both its ancestor and descendant enzyme do so. X, xanthine; XR, xanthosine; 1X, 1-methylxanthine; 3X, 3-methylxanthine; 7X, 7-methylxanthine; TP, theophylline; TB, theobromine; PX, paraxanthine.

After gene duplication of AncIpCS1, one daughter enzyme ultimately gave rise to IpCS1, which exhibits preference to methylate xanthine to produce 3X (Fig. 5a). The other daughter enzyme, AncIpCS2, appears to have maintained highest activity with X, 3X and 7X like AncIpCS1 (Fig. 5a). However, unlike its ancestor, AncIpCS1, AncIpCS2 evolved high relative activity with 7X to produce not just TB, but also PX by methylation at the N1 position (Fig. 5a and Supplementary Fig. 6b). AncIpCS2 retained the ancestral activity of AncIpCS1 with xanthine to produce 1X, but also evolved the ability to methylate X at the N7 position (Fig. 5b and Supplementary Fig. 6). This enzyme also retained ancestral activity with 3X to produce only TB by N7 methylation but lost the ability to methylate 3X at the N1 position to form TP. These activities of AncIpCS2 would have allowed for ancestral flux to produce 1X, 7X, TB and PX but not caffeine. Because a YM ancestor could have possessed both AncIpCS2 and a descendant of AncIpCS1, AncIpCS1’ (Fig. 5b), additional pathway flux is possible. If AncIpCS1’ retained activities of its ancestor, AncIpCS1, then the ancestral *Ilex* species could have also produced 3X and TP making for an even more diverse array of xanthine alkaloids in its tissues (Fig. 5b). It has been shown that the xanthine alkaloids, 1X, 3X, and TP, can bind to modern-day rat adenosine receptors^48^. Therefore, if these molecules were to accumulate in ancestral *Ilex* tissues, they could have conferred a selective advantage which would likely result in retention of the ancient genes. Ultimately, once gene duplication led to the generation of the three modern-day CS-type enzymes in YM, pathway flux could be channelled away from intermediates like 1X and TP such that the modern-day pathway to caffeine evolved (Fig. 5b). Not only did the modern-day CS enzymes of YM evolve to catalyse a pathway from X>3X>TB>CF from ancestral biosynthetic networks of different products, *Theobroma* and *Paullinia* also independently evolved enzymes with similar properties^19^. And, they did so from ancestral pathways that, like YM, had alternative ancestral fluxes^24^. While it could be due to chance alone that all three lineages converged to catalyse a similar pathway from differing ancestral networks, it is also possible that it was advantageous to specialize for flux to TB via X and 3X because either it is more enzymatically favourable or these intermediates have greater adaptive value than other structural isomers.

### Protein crystal structure of IpCS3 reveals convergent structural basis for methylation of theobromine to form caffeine

We successfully crystallized and determined the 2.7 Å resolution structure of IpCS3 (PDB ID: 8UZD), that converts TB into CF. This enzyme crystallizes as a holo-homodimer, bound to both of its reaction products: S-adenosyl-homocysteine (SAH) and caffeine (Fig. 6a, Supplementary Table 8). As is typical for the SABATH family of methyltransferases, IpCS3 exhibits a Rossman-like fold composed of 7 β-strands surrounded by 5 α-helices which bind the methyl-donor S-adenosyl-L-methionine (SAM), as well as an alpha-helical cap which bind the methyl-acceptor theobromine^49–52^. This structural information of the enzyme bound to both of its products, SAH and caffeine, facilitates an in-depth comparison of the active site structures of the caffeine-producing CS-type enzyme found in *Ilex* and the XMT-type enzyme in *Coffea*^49^ to determine the extent to which convergence of physicochemical properties of the active site has allowed for independent specialization for theobromine methylation by the paralogous SABATH enzymes. Although the IpCS3 structure was obtained in complex with its product, caffeine, it can be assumed that the binding mode is conserved for its precursor, theobromine (Supplementary Fig. 7). Indeed, our computational modelling of theobromine in the active site of IpCS3 predicts it to be oriented as we have discerned from the diffraction data (Supplementary Fig. 8). Thus, in the following comparisons, the atomic numbering for the theobromine precursor will be used to facilitate comparison to the DXMT structure.

**Fig 6.**
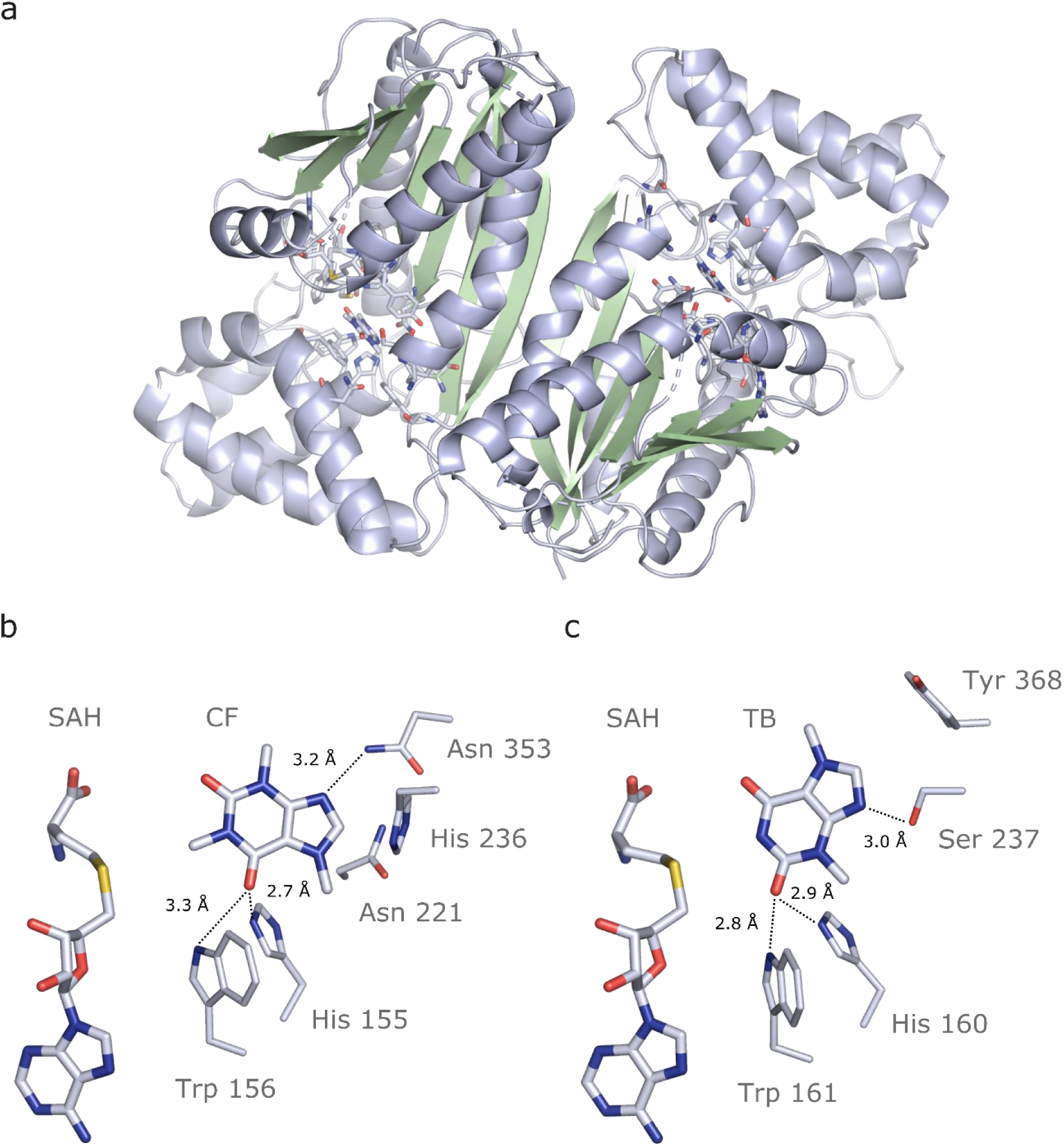
Crystal structure of IpCS3 in complex with Caffeine (CF) and S-adenosyl-homocysteine (SAH) and comparison with the active site of *Coffea* DXMT. **a,** Overview of the crystal structure of IpCS3 (PDB ID: 8UZD) depicting the active site of the enzyme in complex with CF and SAH. **b**, Relevant residues in IpCS3 for ligand recognition. **c**, Relevant residues in DMXT (PDB ID: 2EFJ) for ligand recognition. Protein residues are displayed as lines with carbon atoms coloured in bluewhite while small molecules - caffeine (CF), theobromine (TB), and S-adenosyl-homocysteine (SAH) - are drawn as sticks. Colour code for the rest of the atoms: nitrogen (blue), oxygen (red) and sulphur (yellow). Hydrogen bond interactions are indicated as black dotted lines.

In both DXMT and IpCS3, there are several conserved residues, shared by nearly all SABATH enzymes, that form part of the active site pocket and appear to play important roles in binding many different substrates^50, 51^. His160 and Trp161 in DXMT are in the same relative positions as His155 and Trp156 in IpCS3 (Fig. 6b, c). These residues are ca. 3 Å from TB and participate in H-bonding but to different atoms of the substrate. In DXMT, the NE1 of Trp161 and NE2 of His160 form hydrogen bonds to carbonyl O2 of TB when positioned for N1 methylation; yet, in the structure of IpCS3 these corresponding side chain groups form hydrogen bonds to O6 of TB. Despite these two residues being conserved for H-bonding, the substrates are rotated 180 degrees along an axis going through N1 and C4. Thus, the conserved His and Trp residues interact with opposing carbonyls in TB/CF but still position the substrate for N1 methylation (Fig. 6b, c).

On the other hand, there are residues that differ between the two enzymes but appear to provide for important substrate interactions. Specifically, in the structure of DXMT, the hydroxyl group of Ser237 allows specific hydrogen bonding with N9 to position TB for N1 methylation (Fig. 6c). In IpCS3, His236 is found at the homologous position in the structure. Nevertheless, its involvement in H-bonding with N9 is uncertain as the distance between nitrogen atoms is ca. 4Å. Tyr368 of DXMT is found to participate in π-π interactions with the ring structure of TB^53^; yet in IpCS3, Asn353 is found in the homologous position and the amine forms a hydrogen bond with N9 due to its proximity within 3.2 Å, which is additionally stabilized by Asn221 and His236 (Fig. 6b). Because the residues in these positions of IpCS3 and DXMT differ yet contribute to TB binding, these independent replacements represent convergent structural solutions for N1 methylation of the substrate.

### Computational modelling of IpCS1 and IpCS2 active sites predict convergent substrate binding residues for xanthine and 3-methylxanthine methylation

Previous studies used site-directed mutagenesis of two sequence regions in CS-type caffeine biosynthetic enzymes from *Theobroma* and *Paullinia* to uncover the mutational basis for the convergent evolution of substrate preference switches towards their preferred substrates, X and 3X^24^. In order to determine whether the same regions were convergently mutated in IpCS1 and IpCS2 and provide binding interactions with X and 3X, respectively, AlphaFold2^54^ models and subsequent docking studies were performed (Supplementary Fig. 9). Modelling of substrate binding in the predicted active sites of IpCS1 and IpCS2 (Fig. 7a, b) shows that the preferred substrates have optimal binding orientations that would result in methylation to form the products that were experimentally detected in our assays shown in Fig. 4. From our docking simulations, IpCS1 residues W156, N221 and Y265 are positioned for hydrogen bonding with the carbonyl moieties of xanthine to position N3 for methyl transfer from SAM (Fig. 7a). Although *Theobroma* and *Paullinia* CS1 enzymes specialized for xanthine methylation also possess W156 and Y265 in the homologous positions (Supplementary Fig. 10), these residues are highly conserved amongst nearly all angiosperm SABATH enzymes. On the other hand, the homologous position to N221 did not change concomitantly with the evolution of X preference in either *Theobroma* and *Paullinia* (Supplementary Fig. 10); instead, when their “region III” was mutated, activity with X improved in both^24^. Because IpCS1 was not mutated in the homologous region III, there appear to be convergent solutions allowing for efficient positioning of X for 3X biosynthesis amongst these enzymes. In the case of IpCS2, two hydrogen bond donors S24 and T25 appear to contribute to the positioning of 3X in the active site (Fig. 7b). This homologous region was experimentally mutated in *Theobroma* and *Paullinia* CS2 enzymes and improved specialization for 3X methylation in both, although the actual substitutions are different in each case^24^. Thus, this may represent an additional instance where convergent mutations of the same region lead to specialization for 3X methylation.

**Fig. 7.**
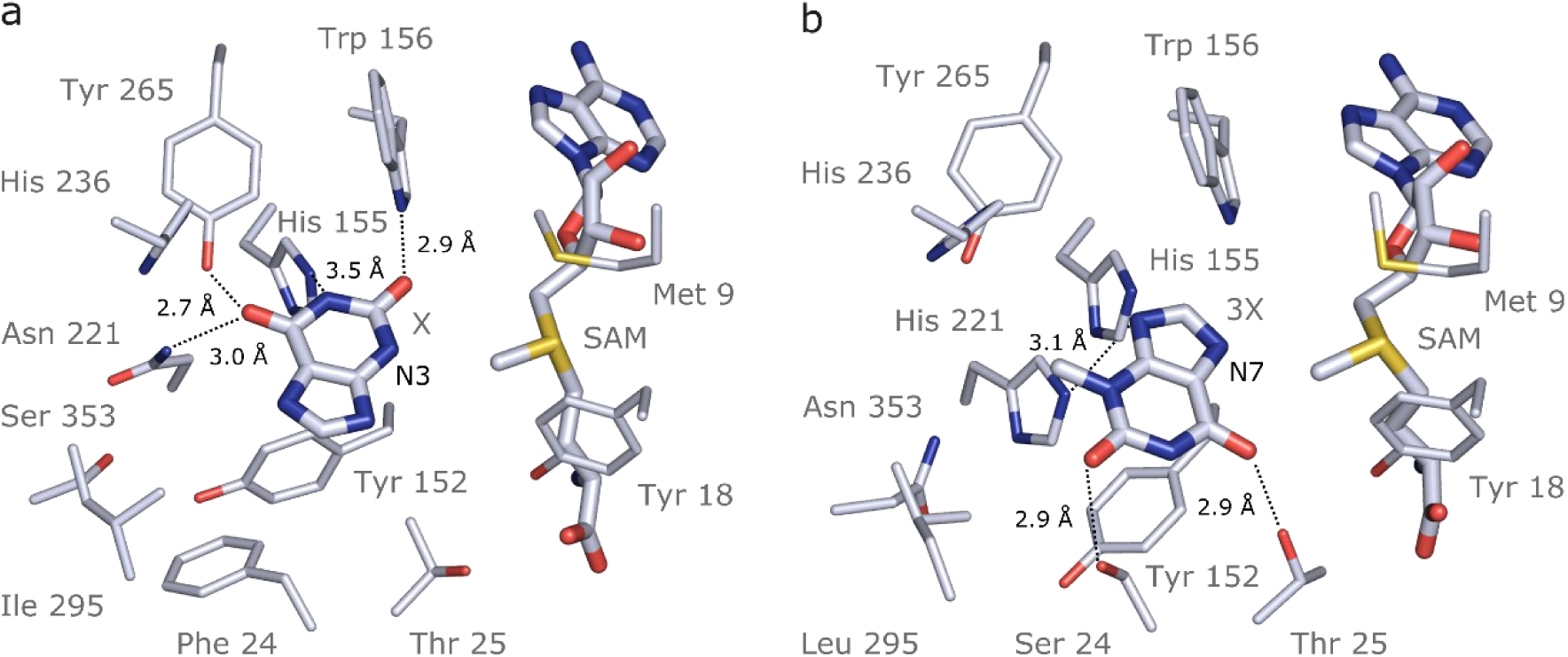
Docking models of xanthine alkaloids in IpCS1 and IpCS2 active sites. **a**, IpCS1- X complex. **b**, IpCS2-3X complex. Protein residues are displayed as lines with carbon atoms coloured in bluewhite while small molecules - xanthine (X), 3-methylxanthine (3X), caffeine (CF), paraxanthine (PX), S-adenosyl-L-methionine (SAM) and S-adenosyl-homocysteine (SAH) - are drawn as sticks. Colour code for the rest of the atoms: nitrogen (blue), oxygen (red) and sulphur (yellow). Hydrogen bond interactions are indicated as black dotted lines.

### Comparative phylogenomic analyses of caffeine biosynthetic genes reveal historical constraints to convergent gene co-option

Many nearly ubiquitous specialized metabolites involved in defence, development and floral scent are produced by SABATH enzyme family members that appear to be conserved across diverse angiosperm lineages, such as SAMT that methylates salicylic acid^55^ and IAMT that methylates Indole-3-acetic acid^52^. However, caffeine is sporadically distributed amongst disparate angiosperm lineages and seems to have only recently evolved by convergence in a few distantly related orders^19^. Our comparative evolutionary genomic analysis of the CS and XMT syntenic regions across angiosperm (Fig. 8) indicates that predicting which SABATH locus a given lineage might co-opt for caffeine biosynthesis is more dependent upon the idiosyncratic history of gene loss than phylogenetic relatedness. For example, in the case of the CS syntenic region used for caffeine biosynthesis in YM and *Theobroma*, *Coffea* lacks a CS ortholog and none can be detected from its genome (Supplementary Fig. 11a). Thus, only XMT was historically available for recruitment in *Coffea*. Conversely, YM appears to have lost any vestiges of XMT orthologues known to be responsible for caffeine biosynthesis in *Coffea* and *Citrus* (Supplementary Fig. 11b-d). This lack of genomic potential may be seen as an evolutionary constraint to gene recruitment for caffeine biosynthesis in *Coffea* and YM.

**Fig. 8.**
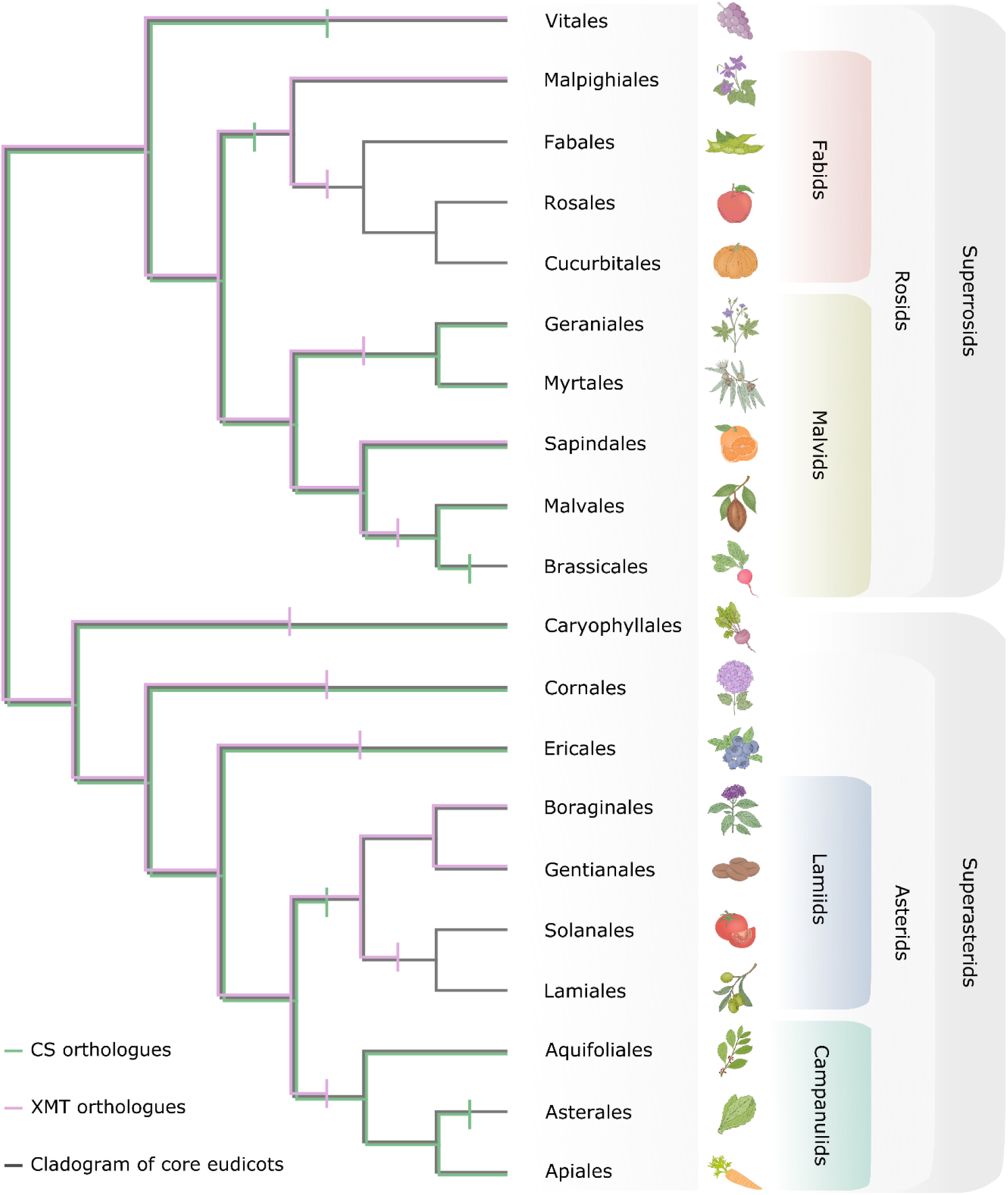
Only CS genes were available for co-option and utilization for xanthine alkaloid biosynthesis in yerba mate whereas coffee only had XMT genes. Both CS- and XMT-type caffeine biosynthetic enzymes were present in the ancestor of core eudicots but numerous apparent losses of one or the other or both has occurred during lineage diversification. Gene loss is represented by vertical bar on relevant branches of the cladogram.

A broader phylogenetic perspective on the XMT and CS syntenic regions provides further insight into genomic canalization and allows for predictions about the underlying genetic basis for caffeine biosynthesis in as-of-yet characterized lineages. As shown in Fig. 8, several angiosperm lineages have neither XMT nor CS and this may explain why caffeine has apparently never evolved in the large and diverse orders Brassicales, Asterales, Solanales and Lamiales even though it has been shown to be advantageous in transgenic plants^56, 57^. In the case of *Cola*, a caffeine-producing genus from Africa^58^, it is predicted to have co-opted CS genes for xanthine alkaloid methylation because the order Malvales to which it belongs appears to have lost XMT orthologues prior to its origin (Fig. 8). Tests of this hypothesis await genomic sequences and functional studies of *Cola* enzymes. However, even with a functional XMT or CS enzyme, duplication and diversification appears to be required to assemble a complete pathway to caffeine as shown here for YM.

## Methods

### Plant materials

*Ilex paraguariensis* A. St.-Hil. var. *paraguariensis*, cv CA 8/74 (INTA-EEA Cerro Azul, Misiones, Argentina) and cv SI-49 (Establecimiento Las Marías S.A.C.I.F.A., Corrientes, Argentina) were used in this study. High productivity, increased tolerance to drought, and ease of propagation with stem cuttings characterize these genotypes^27, 59, 60^.

### DNA extraction and sequencing

Two DNA extraction and sequencing approaches were combined to improve the accuracy of genome assembly. First, young leaves of cv CA 8/74, preserved in silica-gel, were used to isolate total genomic DNA with the DNeasy® Plant Mini Kit (QIAGEN®), following manufacturer’s instructions. Paired-end libraries (with insert sizes of 350 bp and 550 bp) and mate-pair libraries (with insert sizes of 3 kbp, 8 kbp and 12 kbp) were constructed using the Illumina TruSeq® DNA Sample Preparation Kit and Illumina Nextera® Mate Pair Library Preparation Kit following the kit’s instructions, respectively. The obtained libraries were sequenced on an Illumina HiSeq 1500 platform, generating ∼263.2 Gb of raw data. Second, young leaves of cv SI-49, preserved in silica-gel, were used to purify high molecular weight DNA with the Quick-DNA™ HMW MagBead Kit (Zymo Research), according to the manufacturer’s instructions. Long-reads libraries were prepared using Sequel Binding Kit 1.0 (Pacific Biosciences), following manufacturer’s instructions. The obtained libraries were subsequently sequenced on PacBio Sequel I (Pacific Biosciences) using Sequel Sequencing Kit 1.0 (Pacific Biosciences) and SMRT Cell 1M (Pacific Biosciences), generating ∼77.5 Gb of additional raw data.

### Genome assembly and quality assessment

We opted for a pipeline that could take advantage of both short and long sequencing technologies. For the short-reads, we applied Trimmomatic^61^ v.0.39 to remove adaptor contaminations and filter low-quality reads (reads with mean quality scores ≤ 25, reads where the quality of the bases at the head or tail was ≤ 10 and reads with a length ≤ 30 bp). The resulting clean reads were then corrected using Quake^62^ v.0.3. Contig assembly and scaffolding was done using the assembler SOAPdenovo^63^ v.2 (55-mer size), with the mate-pair reads being used to link contigs into scaffolds. After the assembly, DeconSeq^64^ v.0.4.3 was used to detect and remove sequence contaminants. Contigs and scaffolds clearly belonging to the chloroplast and mitochondria genomes were also discarded. YM transcriptome sequences^25–27^ and public databases (KOG^30^ and DEG^31^) were used to validate the genome assembly. Canu^65^ v.2.2 was used to perform the self-correction and assembly of the long reads, using the default parameters and stopOnLowCoverage=20. For both short and long assemblies, we separated the assembly haplotypes (haplotigs) using PurgeHaplotigs^66^ with the recommended parameter values. Then, we merged both SOAPdenovo v.2 and Canu v.2.2 curated assemblies using Quickmerge^67^ v.03, where only contigs with minimum overlap of 5000 bp (-ml 5000) were merged and only the contigs greater than 1000 bp (-l 1000) were retained. The resulting scaffolds and contigs were refined further with the gap-filling module in SOAPdenovo v.2 (GapCloser) and SSPACE^68^ v.2.1.1.

### Gene prediction and annotation

First, we masked the genome assembly with RepeatMasker (http://repeatmasker.org/). Then, we predicted the protein coding and non-coding genes using Funannotate^69^ v.1.8.13 previously training it with the available *Ilex paraguariensis* RNA-Seq experiments (NCBI projects PRJNA315513, PRJNA375923 and PRJNA251985). Then, Infernal^70^ v.1.1.4 was employed to improve the prediction of small RNAs and microRNAs, while tRNAScan-SE^71^ v.2.0 was used to improve the prediction of transfer RNAs. Ribosomal RNAs were predicted using RNAmmer^72^ v.1.2. The TAPIR web server^73^ (http://bioinformatics.psb.ugent.be/webtools/tapir) and the TargetFinder software^74^ v.1.7 were used to identify miRNA targets. InterProScan^75^ v.5.55-88.0 and eggNOG-mapper^76^ v.2.1.7 were employed for the functional assignment of the predicted genes.

### Repeat content estimation

The repeat content was estimated employing Dfam TE Tools v.1.5 (https://github.com/Dfam-consortium/TETools). First, we used RepeatModeler v.2.0.3^77^ to build a database with *Ilex* repeat families. Then, we merged that database with Dfam v3.6^78^ and GIRI Repbase ver 20181026^79^. Finally, we ran RepeatMasker on the assembly using the merged database to look for repeat sequences.

### Genome duplication analysis

Rates of synonymous substitution (K_s_) between paralogous genes and orthologous genes in *Lactuca sativa*, *Daucus carota*, *Ilex paraguariensis*, *Coffea canephora* and *Vitis vinifera* were determined using CoGe’s tool SynMap (https://genomevolution.org/). Gaussian mixture models (GMMs) were fitted to the resulting K_s_ distributions with the mclust R package^80^ v.5.0, and significant peaks were identified using the SiZer R package^81^ v.0.1-7. To estimate the age of the lineage-specific polyploidization event (Ip-α) in *Ilex*, we considered two different phylogenies (a multiple nuclear genome phylogeny and a plastid genome phylogeny). With the median K_s_ value of yerba mate-grape orthologues (∼0.89) and the divergence date of the two species (125.64 Ma for the multiple nuclear genome phylogeny and 123.7 Ma for the plastid genome phylogeny), we calculated the number of substitution per synonymous site per year (*r*) for YM (divergence date = *K_s_*/(2 x *r*)). Conforming to the multiple nuclear genome phylogeny, the YM *r* value is 3.54E-9; while for the plastid genome phylogeny, the YM *r* value is 3.59E-9. These *r* values and the SiZer K_s_ range of YM paralogues (∼0.35-0.5) were then applied to estimate the age of Ip-α. Finally, to determine the syntenic depth ratio between *I. paraguariensis* and *C. canephora* and *V. vinifera*, we employed CoGe’s tool SynFind (https://genomevolution.org/), using a distance cutoff of 10 genes and requiring at least 5 gene pairs per synteny block.

### Gene expression quantitation

First, YM transcriptome reads (PRJNA315513) were mapped to IpCS1-5 transcripts, obtained from the *de novo* transcriptome assembly and annotation, using BWA^82^. Then, with the number of mapped reads, the abundance of each transcript was calculated, normalized by transcript length and transcriptome size (quantification in RPKM, reads per kilobase per million mapped reads).

### Cloning, heterologous expression and purification of enzymes

Two different approaches were used to clone IpCS genes: RT-PCR from leaf tissue and custom gene synthesis. For RT-PCR of IpCS2, cDNA was obtained from 1 µg of RNA from fresh YM leaves using standard procedures and cycling conditions with the following two primers: IpCS2F 5’- ATGGACGTGAAGGAAGCAC-3’ and IpCS2R 5’-CTATCCCATGGTCCTGCTAAG-3’. Following amplification, cDNA was cloned using the pTrcHis TOPO® TA Expression Kit (Invitrogen, Carlsbad, CA). Ligation of cDNA into the pTrcHis vector and subsequent transformation into Top10 *E. coli* cells was carried out according to the manufacturer’s protocol. The transformation mixture was incubated overnight at 37°C on LB plates containing 50 µg/mL ampicillin. Colonies were screened by PCR to obtain full-length inserts that were subsequently verified for insert orientation by DNA sequencing. For IpCS1 & 3 and AncIpCS1 & 2, gene sequences were synthesized by GenScript with codons optimized for expression in *E. coli*. Synthesized genes were subcloned from the pUC57 cloning vector into the pET-15b (Novagen) expression vector using 1.5 μg of DNA and NdeI and BamHI in 30 μl reactions. Linear fragments corresponding to the expected sizes were gel purified using the QIAEX II Gel Extraction Kit (Qiagen Corp.) according to the manufacturer’s instructions. Purified DNA fragments were ligated into pET-15b using T4 DNA ligase from New England Biolabs. Ligation products were transformed into Top10 *E. coli* cells using 2 μl of the ligation reaction. Minipreps of positive transformants were obtained using a QIAprep Spin Miniprep Kit (Qiagen Corp.) and 10 ng of each plasmid was used to transform BL21 *E. coli* cells using standard plating and incubation methods.

Induction of His_6_-protein was achieved in 50 ml cell cultures of BL21 (DE3) with IpCS1 & 3 and AncIpCS1 & 2 in pET-15b or Top10 with IpCS2 in pTrcHis with the addition of 1mM IPTG at 23° C for 6 hours. Purification of the His_6_-tagged protein was achieved by TALON® spin columns (Takara Bio) following the manufacturer’s instructions. Bradford assays were used to determine purified protein concentration, and recombinant protein purity was evaluated on SDS-PAGE gels.

### Enzyme assays

All enzymes were tested for activity with the eight xanthine alkaloid substrates shown in Fig. 1. Radiochemical assays were performed at 24° C for 60 minutes in 50 µl reactions that included 50 mM Tris-HCl buffer, 0.01 µCi (0.5 µl) ^14^C-labelled SAM, 10-20 µl purified protein and 1 mM methyl acceptor substrate dissolved in 0.5 M NaOH. Negative controls were composed of the same reagents, except that the methyl acceptor substrate was omitted and 1 µl of 0.5 M NaOH was added instead. Methylated products were extracted in 200 µl ethyl acetate and quantified using a liquid scintillation counter. The highest enzyme activity reached with a specific substrate was set to 1.0 and relative activities with remaining substrates were calculated. Each assay was run at least twice so that mean, plus standard deviation, could be calculated.

### High performance liquid chromatography (HPLC)

Product identity of enzyme assays was determined using HPLC on 500 μl scaled-up reactions utilizing all the same reagents as described above except that non-radioactive SAM was used as the methyl donor and reactions were allowed to progress for 4 hours. Whole reactions were filtered through Vivaspin® columns (Sartorius) to remove all protein prior to injection in the HPLC. Mixtures were separated by HPLC using a two-solvent system with a 250 mm x 4.6 mm Kinetex® 5 μM EVO C18 column (Phenomenex). Solvent A was 99.9% water with 0.1% TFA and Solvent B was 80% acetonitrile, 19.9% water and 0.1% TFA and a 0-20% gradient was generated over 16 minutes with a flow rate of 1.0 ml/minute. Product identity was determined by comparing retention times and absorbance at 254 and 272 nm of authentic standards. Reactions were compared to negative controls in which no methyl acceptor substrates were added.

### Phylogenetic analyses

In order to accurately determine the orthology of YM SABATH sequences encoded in the genome, we compared them to all previously characterized gene family members in other species. We also included CS and XMT orthologues from relatives from the orders of caffeine-producing species (Malvales, Ericales, Gentianales, Sapindales) available in public databases (GenBank, OneKP) as shown in Fig. 3. Accession numbers for all sequences are provided in Supplementary Table 9. Alignment of amino acid sequences was achieved using MAFFT^83^ v.7.0 and employing the auto search strategy to maximize accuracy and speed. A phylogenetic estimate was obtained using FastTree^84^ v.2 assuming the Jones-Taylor-Thorton model of amino acid substitution with a CAT approximation using 20 rate categories. Reliability of individual nodes was estimated from local support values using the Shimodaira-Hasegawa test as implemented in FastTree.

### Ancestral sequence resurrection

In order to obtain accurate ancestral CS protein estimates, we assembled two datasets to assess variation in terms of sampling. The first dataset included 154 sequences including all CS-type enzymes we could retrieve from GenBank and China National Gene Bank as well as representatives of all other functionally characterized clades of SABATH enzymes (Supplementary Fig. 2). This set of ancestral enzymes was obtained using only the sequences from YM which were available during our initial analyses. Subsequently, once additional *Ilex* genomes became available, we estimated a second set of ancestral sequences using 29 CS-type enzymes from asterids to assess uncertainty in our estimates (Supplementary Fig. 3). IQTree^85^ was used to obtain trees describing the relationships amongst the sequences for both datasets. For the first, the Jones, Taylor, and Thorton (JTT) matrix model for amino acid substitution and the Free rate model for among-site rate heterogeneity^86^ was determined to be the best fit. For the second dataset, the Q matrix as estimated for plants^87^ with a gamma model for rate heterogeneity was the preferred model. Confidence in ancestral sequences was high in both analyses as indicated by the average site-specific posterior probability for AncIpCS1 (0.99) and AncIpCS2 (0.99). In order to determine ancestral protein lengths in regions with alignment gaps, we coded each gap for the number of amino acids possessed and used parsimony to determine ancestral residue numbers as in our previous studies^19^. The estimated sequences were synthesized by Genscript Corp. and had codons chosen for optimal protein expression in *Escherichia coli*. Although the two separate estimates are highly similar to one another (>95% in both cases), the two AncIpCS1 proteins differ at 10 positions and those for AncIpCS2 differ at 7 positions (Supplementary Fig. 4 & 5).

### Crystallization, data collection, phasing and refinement of IpCS3

Initial crystallization screening was performed using the IpCS3 methyltransferase at a concentration of 30 mg/mL incubated with 2 mM TB and 2 mM SAM. Sitting-drop for crystallization screening was set up by equal volume of precipitant and protein solution (0.25:0.25 µL) using a Crystal Gryphon robot (Art Robbins Instruments) and a reservoir volume of 45 µL. Trays were incubated at 9 ℃. Initial hits were further optimized using the hanging-drop method at 9°C, with 150 µL reservoir solution and 1:1 ratio of precipitant to protein and ligand solution in a 2 µL drop. Attempts to crystallize with SAH or uncleavable SAM analogs and TB to attain a pre methylation structure were unsuccessful given the poor diffraction of these crystals. Therefore, the latter was composed of 33 mg/mL IpCS3 protein concentration, 4 mM TB and 2 mM SAM, and the crystallization condition was optimized to 25% PEG 3350, 0.2 M NH_4_SO_4_, 0.1 M Bis-Tris methane pH 5.5. Square crystals grew over 10 days, but initial X-ray crystallography data revealed a poor electron density for SAH and an electron density in the active site for CF, the product, rather than for TB. Consequently, crystals were grown in the aforementioned condition and subsequently soaked for 4h at 9°C in the precipitant solution supplemented with 10 mM SAH and 10 mM TB. The idea was to supply an excess of the expended methyl source and additional TB to convert any existing SAM as we did not have access to caffeine as a reagent. Crystals were transiently soaked in the precipitant solution supplemented with 20% Ethylene Glycol immediately prior to vitrification by direct immersion into liquid nitrogen. Diffraction data were collected at the Advanced Photon Source (APS) at Argonne National Laboratory Sector-21 via the Life Sciences-Collaborative Access Team (LS-CAT) at beamline 21-ID-G. Diffraction data were indexed, integrated, and scaled using the autoPROC software package^88^. The structure was solved by molecular replacement using Phaser-MR included in the Phenix software package^89^, using PDB ID 6LYH structure as the replacement model. The model was subject to rounds of manual building followed by refinement using REFMAC5^90^, and was manually built in COOT^91^ v.0.9.8.3. Crystallographic statistics are listed in Supplementary Table 8.

### Structure prediction and molecular docking

Protein structures of IpCS enzymes were predicted using the ColabFold implementation of AlphaFold2^54^ with no template. Diagnostic plots depicting the MSA coverage, alignment error and LDDT are shown in the supplementary information (Supplementary Fig. 9). Structures of xanthine alkaloid ligands (X, 3X, and TB) were downloaded from the ChEMBL database^92^; protonation states were checked by Chemicalize^93^ and optimized using the VMD Molefacture plugin^94^. The receptor structures were prepared following the standard AutoDock protocol^95^ using the prepare_receptor4.py script from AutoDock Tools. All non-polar hydrogens were merged, and Gasteiger charges and atom types were added. The ligand PDBQT was prepared using the prepare_ligand4.py script. The grid size and position were chosen to contain the whole ligand-binding site (including all protein atoms closer than 5 Å from all ligands). For each system, 10 different docking runs were performed. Docking was performed using AutoDock Vina v.1.2.0^96^. The docking results were further analysed by visual inspection. Images of the molecules were prepared using the PyMOL molecular graphics system^97^.

### Synteny comparisons and phylogenetic distribution of CS and XMT

The presence or absence of CS and XMT genes was determined for orders of plants for which at least one genomic sequence exists, as shown in Fig. 8. For those species which do not yet have an available assembly, we used BLAST^98^ analyses of GenBank (nr and TSA databases), Phytozome^99^ as well as the OneKP dataset^37^ in China National GeneBank. BLAST combined with subsequent phylogenetic analyses were also used to verify presence/absence of CS- or XMT-type sequences in cases where the syntenic regions did not appear to encode one or the other gene. Comparisons of the CS and XMT syntenic regions were performed using CoGe’s tool GEvo (https://genomevolution.org/).

## Data availability

The Illumina and PacBio raw sequence data, assembly and annotation were deposited in the European Nucleotide Archive (ENA) under BioProject No. PRJEB65927. An assembly obtained only with the Illumina data was also deposited in ENA under BioProject No. PRJEB36685. The plasmids used to produce proteins are freely available upon request. The atomic coordinates and structure factors have been deposited in the Protein Data Bank, Research Collaboratory for Structural Bioinformatics, Rutgers University, New Brunswick, NJ (http://www.rscb.org) with the accession code 8UZD for the IpCS3 structure bound to caffeine and SAH.

## Acknowledgments

This work was supported by Consejo Nacional de Investigaciones Científicas y Técnicas de Argentina (CONICET); PRO.MATE.AR project, funded by Secretaría de Políticas Universitarias del Ministerio de Educación de la Nación Argentina; CABANA project, funded by UKRI-BBSRC on behalf of the Global Challenges Research Fund (BB/P027849/1), the U.S. National Science Foundation (NSF) (grants MCB-1120624 and MCB-2325341 to T.J.B.); and the European Molecular Biology Laboratory (EMBL). We would like to express our gratitude to Centro de Cómputos de Alto Rendimiento (CeCAR) and UBA-FCEN-QB-Cluster for providing access to computational resources, which facilitated the majority of computational analyses in this work. Special thanks are extended to Kevin Blair and the Department of Chemistry for facilitating our HPLC analyses.

## Contributions

F.A.V., C.P.M., T.J.B. and A.G.T. designed the project and coordinated the research activities. F.A.V., T.J.B. and A.G.T. wrote the manuscript with input from all co-authors. F.A.V. was involved in all analyses. R.M.A. extracted the genomic DNA. E.J.S., G.F.B., R.R.M.O., G.L.N., R.M.A., and G.O. performed the assembly and annotation of the genome. F.A.V. conducted the evolutionary and syntenic-gene analyses. A.H.G. and T.J.B. carried out the expression, purification and activity assays of CS enzymes. L.A.D made the AlphaFold2 models. L.A.D. and C.P.M. performed the molecular dockings. A.H.G. and S.N. obtained the crystal structure of IpCS3-CF complex. T.J.B., M.R., and P.D.Z. performed the comparative genomic analyses of caffeine-producing enzymes across angiosperms. P.D.Z., D.A.M., P.A.S., G.O., S.N., T.J.B., and A.G.T. obtained funding to carry out the project. All authors read and approved the final article.

## Competing interests

The authors declare no competing interests.

## Notes

https://www.ebi.ac.uk/ena/browser/view/PRJEB65927

https://www.ebi.ac.uk/ena/browser/view/PRJEB36685

### Competing Interest Statement

The authors have declared no competing interest.

### Summary of Updates

Updates have been made to Figures 3 and 5 and the following sections of the Results: "The caffeine biosynthetic pathway in YM evolved from ancestral networks with different inferred flux" and "Protein crystal structure of IpCS3 reveals convergent structural basis for methylation of theobromine to form caffeine". As well as the Method sections: "Plant materials", "DNA extraction and sequencing", "Gene expression quantitation", "Ancestral sequence resurrection", and "Crystallization, data collection, phasing and refinement of IpCS3". In addition, the following have also been revised: data availability, contributions, and Supplemental files.

