## Supplementary Information for "Yerba mate (*Ilex paraguariensis*) genome provides new insights into convergent evolution of caffeine biosynthesis"

### Supplementary Notes

### S1. Non-coding RNAs in yerba mate

#### S1.1. Transfer RNAs

Analysis of transfer RNA (tRNA) family members revealed the presence of 815 tRNA genes including 726 standard tRNAs, 76 pseudo tRNAs, 11 tRNAs with undetermined isotypes and 2 possible suppressor tRNAs (Supplementary Table 4). No selenocysteine tRNA was found in the yerba mate genome, which is consistent with the results of other tRNA analyses carried out in higher plants[1]. According to evolutionary analyses, selenoproteins and selenocysteine insertion sequence (SECIS) elements present in protozoans and animals evolved early, and were independently lost in higher plants and fungi through evolution[2]. With regard to the nonsense suppressor tRNAs, only ochre and opal nonsense suppressor tRNA genes were found in the yerba mate genome, which suppress the phenotypes of ochre and opal mutations respectively.

The length of the standard tRNA sequences ranged from 61 to 236 nucleotides, encoding 53 different anti-codons/isoacceptors in total, with $tRNA^{Ser}$ having the highest abundance of genes and $tRNA^{Tyr}$ having the lowest (Supplementary Table 4). In our study, we also found that yerba mate tRNAs contain introns in 13 of the 55 $tRNA^{Met}$, 9 of the 17 $tRNA^{Tyr}$, 1 of the 65 $tRNA^{Ser}$, and 1 of the 22 $tRNA^{Ile}$ genes, providing additional demonstration that $tRNA^{Met}$ and $tRNA^{Tyr}$ are not the only intron containing tRNAs in the plant kingdom as it was previously believed[3]. The length of these introns varied from 6 to 163 nucleotides and all of them were found at the canonical position, one nucleotide 3′ to the anti-codon loop.

### S1.2. Ribosomal RNAs

Analysis of ribosomal RNA (rRNA) family members showed the presence of 425 5S rRNA, 23 18S rRNA and 23 25S rRNA genes. The large variability in the copy number of the rRNA genes has been observed and studied in plants for decades[4]. It is believed that a high copy number of these genes is important to ensure increased demand of proteosynthesis during plant development, but also to stabilize the cell nucleus[5]. With regard to its genomic organization, the 18S and 25S rRNA genes were clustered together, while the 5S rRNA genes were tandemly located elsewhere in the genome (S-type arrangement). However, some 5S rRNA genes were linked with the 18S and 25S rRNA genes as well (L-type arrangement). Given this observation, we should incorporate the yerba mate genome to the list of eukaryotic genomes with an L-arrangement of ribosomal DNA.

### S1.3. Small RNAs

Analysis of small RNA family members revealed the presence of 348 small nuclear RNA (snRNA) genes including 65 U1 snRNA, 46 U2 snRNA, 29 U4 snRNA, 33 U5 snRNA and 146 U6 snRNA genes corresponding to the major spliceosome complex, and 19 U6atac snRNA, 7 U11 snRNA and 1 U12 snRNA genes corresponding to the minor spliceosome complex. Furthermore, it showed the presence of 2,670 small nucleolar RNA (snoRNA) genes, of which 2,631 (~98.54%) were box C/D snoRNA genes and 39 (~1.46%) were box H/ACA snoRNA genes. Both groups of snoRNAs are involved in the cleavage of precursor ribosomal RNA (pre-rRNA) and determine site-specific modification in pre-rRNAs and snRNAs, though the box C/D snoRNAs are usually associated with 2′-O-ribose methylation, while the box H/ACA snoRNAs are normally associated with 2′-O-ribose pseudouridylation[6,7]. The greater abundance of the box C/D snoRNAs in the yerba mate genome could be explained, first, by the fact that plants have higher numbers of 2′-O-ribose methylated nucleotides than archaea, yeast and other higher eukaryotes, and second, by the fact that computer algorithms still find difficult to predict the relatively short conserved sequences of box H/ACA snoRNAs[7]. It was remarkable the high copy number (2,377) of snoRNA R71 in the yerba mate genome, which is a member of the box C/D family. This snoRNA, which is thought to function as a 2'-O-ribose methylation guide for 18S rRNA, has been identified in multiple copies in most eudicot genomes[8].

S1.4. Micro RNAs

Analysis of micro RNA (miRNA) family members revealed the presence of 226 miRNA genes belonging to 30 families. The most abundant miRNAs usually involved in growth and development were miR156, miR166 and miR159, while the most abundant miRNAs normally involved in stress responses were miR169_2, miR167_1, miR399, and miR395 (Supplementary Table 5). Nevertheless, the functions of miRNAs slightly differ among plants. Therefore, to better understand the regulatory effect of miRNAs in yerba mate, we used the TAPIR web server[9] and the TargetFinder software[10] to identify yerba mate miRNA targets (Supplementary Table 6). The results obtained allowed us to infer the role of 14 of the 30 miRNA families found in the yerba mate genome. Apparently, miR159, miR164, miR169_2 and miR169_5 regulate a variety of processes related to development and stress responses. On the one hand, both miR159 and miR164 regulate the expression of myb-like transcription factors, involved in auxin homeostasis, lateral root and leaf development, leaf senescence, response to abscisic acid, response to the absence of light and response to salt stress. miR164 also regulates the synthesis of vitamin B5 involved in embryo development. On the other hand, both miR169_2 and miR169_5 regulate the expression of a galactinol synthase involved in the response to cold stress, heat stress, oxidative stress, salt stress and water deprivation; a kinesin-like protein involved in pollen development and a mitogen-activated protein kinase involved in directing cellular responses to mitogens, osmotic stress, heat shock and proinflammatory cytokines. miR167_1 is probably involved only in plant growth and development as it regulates the expression of an auxin response factor (arf). And last, miR171_1 and miR390 are likely involved only in stress responses. miR171_1 regulates the expression of an RNA-binding family protein and an endoglucanase involved in host defence, whereas miR390 regulates the expression of a rotamase FKBP 1 involved in the response to heat stress, osmotic stress and wounding. It is important to mention that the functions of all the predicted targets were gathered from the Arabidopsis Information Resource (TAIR)[11], and therefore the functional involvement of these miRNAs in yerba mate must be experimentally validated.

### Supplementary Figures

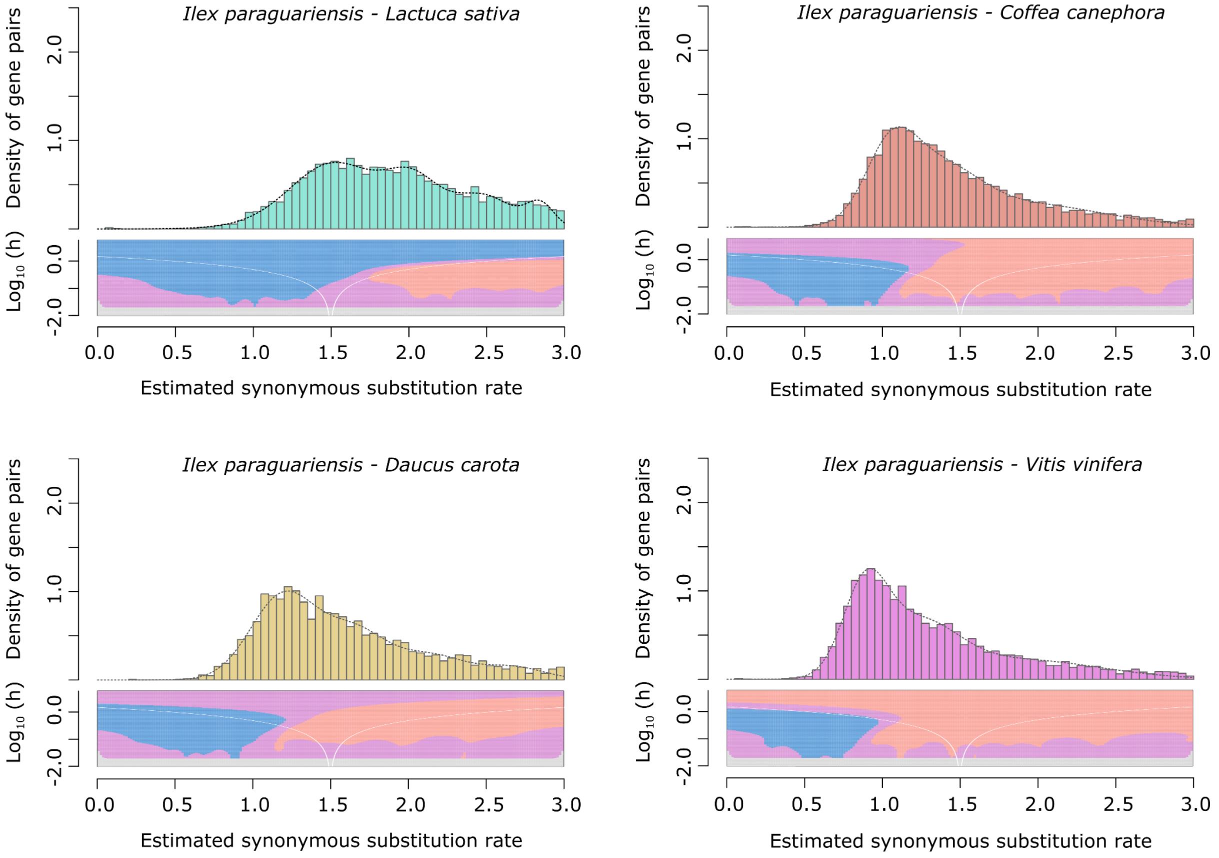

**Supplementary Figure 1. Ks distributions with Gaussian mixture model and SiZer analyses of *I. paraguariensis* and *L. sativa* (green), *D. carota* (yellow), *C. canephora* (red) and *V. vinifera* (purple) orthologues**. SiZer maps below histograms identify significant peaks at corresponding Ks values. Blue represents significant increases in slope, red indicates significant decreases, purple represents no significant slope change, and grey indicates not enough data for the test.

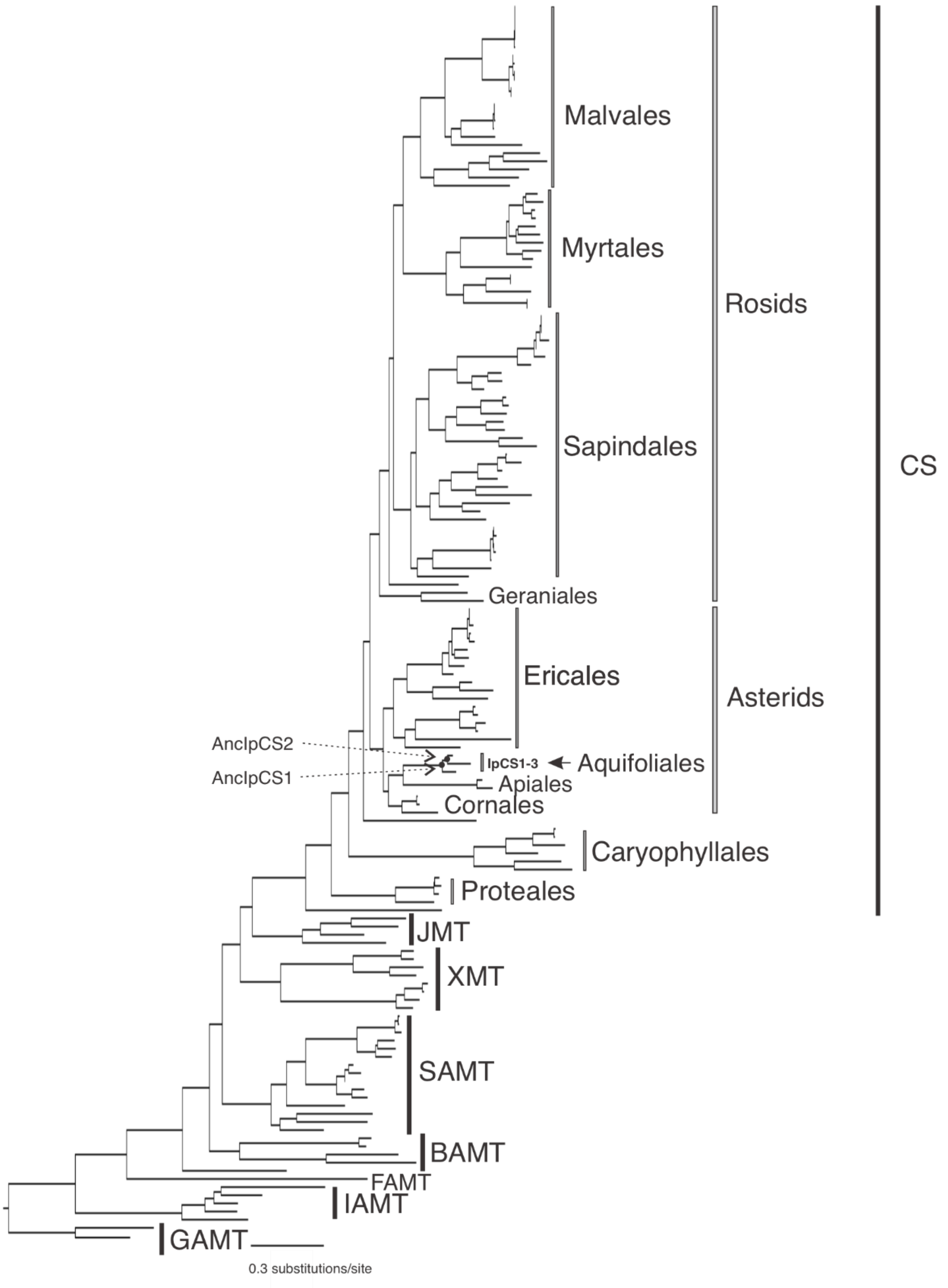

**Supplementary Figure 2. SABATH enzyme family phylogenetic tree used for obtaining ancestral sequence estimates for the clade including IpCS1-3 of Aquifoliales (log-likelihood= -46631.672).** Clades of enzymes for which at least one sequence has been functionally characterized are labeled. GAMT, gibberellin MT; IAMT, indole-3-acetic acid MT; FAMT, farnesoic acid MT; BAMT, benzoic acid MT; XMT, xanthine alkaloid MT used for caffeine biosynthesis in *Coffea* and *Citrus*; SAMT, salicylic acid MT; JMT, jasmonic acid MT; CS, caffeine synthase in *Theobroma*, *Camellia* and *Paullinia*. Within the CS clade, the orders of rosids and asterids are labeled to show interrelationships.

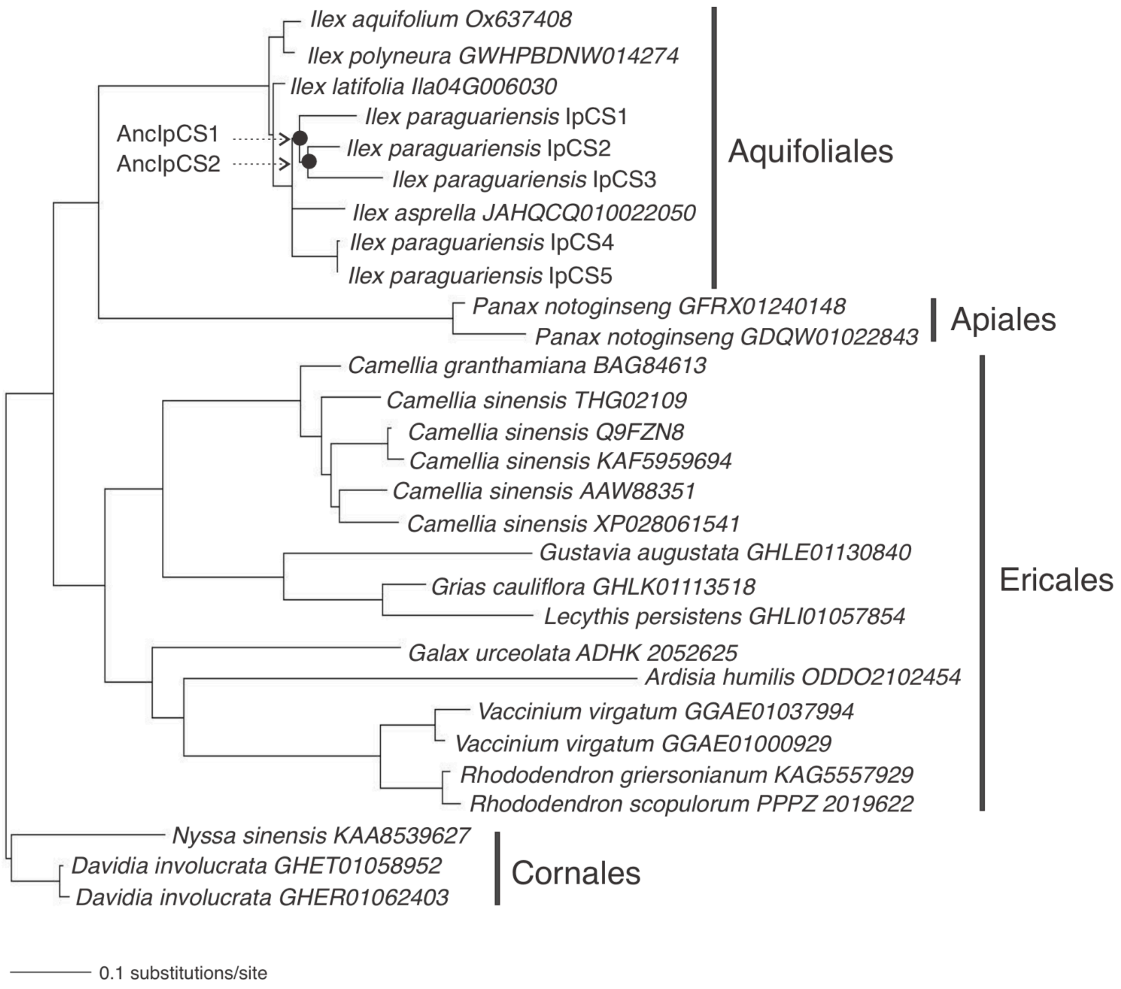

**Supplementary Figure 3. Caffeine synthase enzyme family phylogenetic tree used for obtaining ancestral sequence estimates for AncIpCS1 & 2 (log-likelihood= -7032.8928).**

```
                      10        20        30        40        50        60
AncIpCS1v1 MDVKEALFMNGGEVESSYAQHARFTQKVTSITKPILVNAVHSLFSEDFHRKKVLNVADLG
           :::::::::::::::::::::: :::::::::::::::::::::::::::::::::::::
AncIpCS1v2 MDVKEALFMNGGEVESSYAQHACFTQKVTSITKPILVNAVHSLFSEDFHRKKVLNVADLG
                      10        20        30        40        50        60

                      70        80        90       100       110       120
AncIpCS1v1 CAAGPNTFSVILTVKESLERKCKELNCQPPELQVYLNDLPGNDFNSLFKDLSRVGEDQKS
           :::::: :::::::::::::::::::::::::::::::::::::::::::::::::::::
AncIpCS1v2 CAAGPNPFSVILTVKESLERKCKELNCQPPELQVYLNDLPGNDFNSLFKDLSRVGEDQKS
                      70        80        90       100       110       120

                     130       140       150       160       170       180
AncIpCS1v1 DVLLPCFVMGAPGSFYGRLFPRSSLHLVHSSYSVHWLSQVPKGLTSKEGLPLNKGKIYIS
           ::::::::::::::::::::::: ::::::::::::::::::::::::::::::::::::
AncIpCS1v2 DVLLPCFVMGAPGSFYGRLFPRSWLHLVHSSYSVHWLSQVPKGLTSKEGLPLNKGKIYIS
                     130       140       150       160       170       180

                     190       200       210       220       230       240
AncIpCS1v1 KTSPPVVAAAYLAQFKEDFTLFLKSRAEEMVQNGRMVLILHGRQASDPWGKESCYHWEIL
           ::::::::::::::::::::::::::::::::::::::::::::::::::::::::::::
AncIpCS1v2 KTSPPVVAAAYLAQFKEDFTLFLKSRAEEMVQNGRMVLILHGRQASDPWGKESCYHWEIL
                     190       200       210       220       230       240

                     250       260       270       280       290       300
AncIpCS1v1 AEAISEMVSQGLVDEEKLDSFNVPYYTPLQEEVQDIVDKEGSFAVEHLETFTLDIVDKQE
           :::::::::::::::::::::::::::::::::::::::::::::::.::::: ..:.::
AncIpCS1v2 AEAISEMVSQGLVDEEKLDSFNVPYYTPLQEEVQDIVDKEGSFAVEHIETFTLALADNQE
                     250       260       270       280       290       300

                     310       320       330       340       350       360
AncIpCS1v1 SDTRAKGEQLAKNIRCFTESIISYQFGKEITEKVYHKLTQIVVKDLANRSPTNTSVVVVL
           :::::::::::::::::::::::::::::::::::::::::::::::.::::::.:::::
AncIpCS1v2 SDTRAKGEQLAKNIRCFTESIISYQFGKEITEKVYHKLTQIVVKDLASRSPTNTTVVVVL
                     310       320       330       340       350       360

AncIpCS1v1 SRTMG
           :::::
AncIpCS1v2 SRTMG
```

**Supplementary Figure 4. Alignment of the two estimated sequences for AncIpCS1 that were biochemically characterized in Figure 5.**

```
                   10        20        30        40        50        60
AncIpCS2v1 MDVKEALFMNGGEVESSYAQHARFTQKVTSITKPILVNAVHSLFSEDFHRKKVLNVADLG
           ::::::::::::::::::::::::: ::::::::::::::::::::::::::::::::::
AncIpCS2v2 MDVKEALFMNGGEVESSYAQHACFTQKVTSITKPILVNAVHSLFSEDFHRKKVLNVADLG
                   10        20        30        40        50        60

                   70        80        90       100       110       120
AncIpCS2v1 CAAGPNPFSVILTVKESLERKCKELNCQPPELQVYLNDLPGNDFNSLFKDLSRVGEDQKS
           ::::::::::::::::::::::::::::::::::::::::::::::::::::::::::::
AncIpCS2v2 CAAGPNPFSVILTVKESLERKCKELNCQPPELQVYLNDLPGNDFNSLFKDLSRVGEDQKS
                   70        80        90       100       110       120

                  130       140       150       160       170       180
AncIpCS2v1 DVLLPCFVMGAPGSFYGRLFPRSSLHLVHSCYSVHWLSQVPKGLTSKEGLPLNKGKIYIS
           :::::::::::::::::::::: :::::::::::::::::::::::::::::::::::::
AncIpCS2v2 DVLLPCFVMGAPGSFYGRLFPRSCLHLVHSCYSVHWLSQVPKGLTSKEGLPLNKGKIYIS
                  130       140       150       160       170       180

                  190       200       210       220       230       240
AncIpCS2v1 KTSPPVVAAAYLAQFKEDFTLLLKSRAEEMVQNGRMVLILHGRQASDPWGKESCYHWEIL
           ::::::::::::::::::::::::::::::::::::::::::::::::::::::::::::
AncIpCS2v2 KTSPPVVAAAYLAQFKEDFTLLLKSRAEEMVQNGRMVLILHGRQASDPWGKESCYHWEIL
                  190       200       210       220       230       240

                  250       260       270       280       290       300
AncIpCS2v1 AEAISEMVSQGLVDEEKLDSFNVPYYTPLQEEVQDIVDKEGSFAVEHIETFTLDLVDKQE
           ::::::::::::::::::::::::::::::::::::::::::::::::::::: :.:.::
AncIpCS2v2 AEAISEMVSQGLVDEEKLDSFNVPYYTPLQEEVQDIVDKEGSFAVEHIETFTLALADNQE
                  250       260       270       280       290       300

                  310       320       330       340       350       360
AncIpCS2v1 SDTRAKGEQLAKNIRCFTESIISYQFGKEITEKVYHKLTQIVVKDMANRSPTNTSVVVVL
           :::::::::::::::::::::::::::::::::::::::::::::::.::::::.:::::
AncIpCS2v2 SDTRAKGEQLAKNIRCFTESIISYQFGKEITEKVYHKLTQIVVKDMASRSPTNTTVVVVL
                  310       320       330       340       350       360

AncIpCS2v1 SRTMG
           :::::
AncIpCS2v2 SRTMG
```

**Supplementary Figure 5. Alignment of the two estimated sequences for AncIpCS2 that were biochemically characterized in Figure 5.**

### a. HPLC traces for AncIpCS1 product formation

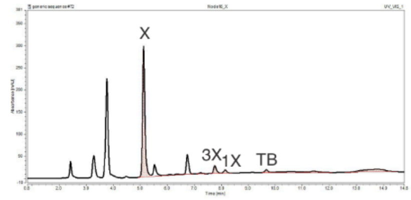

AncIpCS1 + Xanthine

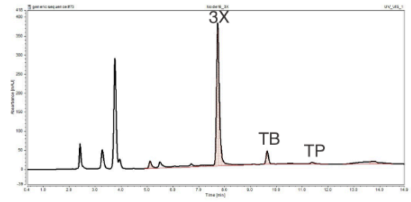

AncIpCS1 + 3-methylxanthine

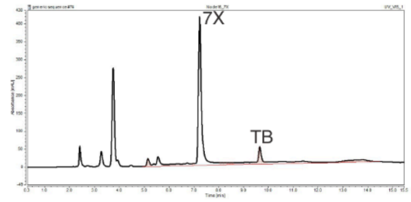

AncIpCS1 + 7-methylxanthine

### b. HPLC traces for AncIpCS2 product formation

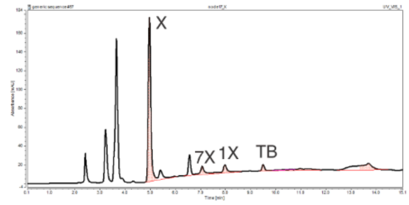

AncIpCS2 + Xanthine

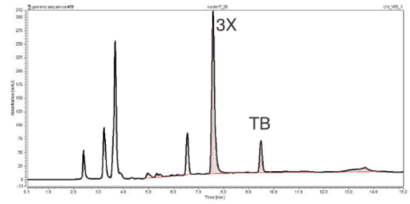

AncIpCS2 + 3-methylxanthine

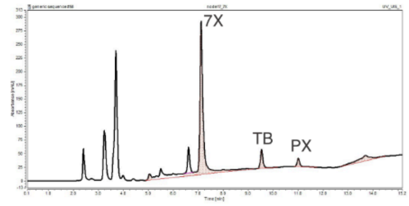

AncIpCS2 + 7-methylxanthine

### c. HPLC trace for xanthine alkaloid standards

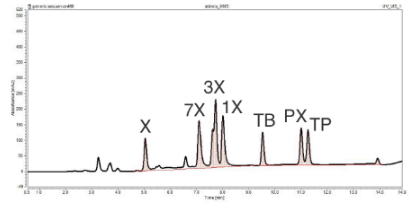

**Supplementary Figure 6. HPLC traces for xanthine alkaloid products formed by ancestral *Ilex* CS enzymes.** X, xanthine; XR, xanthosine; 1X, 1-methylxanthine; 3X, 3-methylxanthine; 7X, 7-methylxanthine; TP, theophylline, TB, theobromine; PX, paraxanthine.

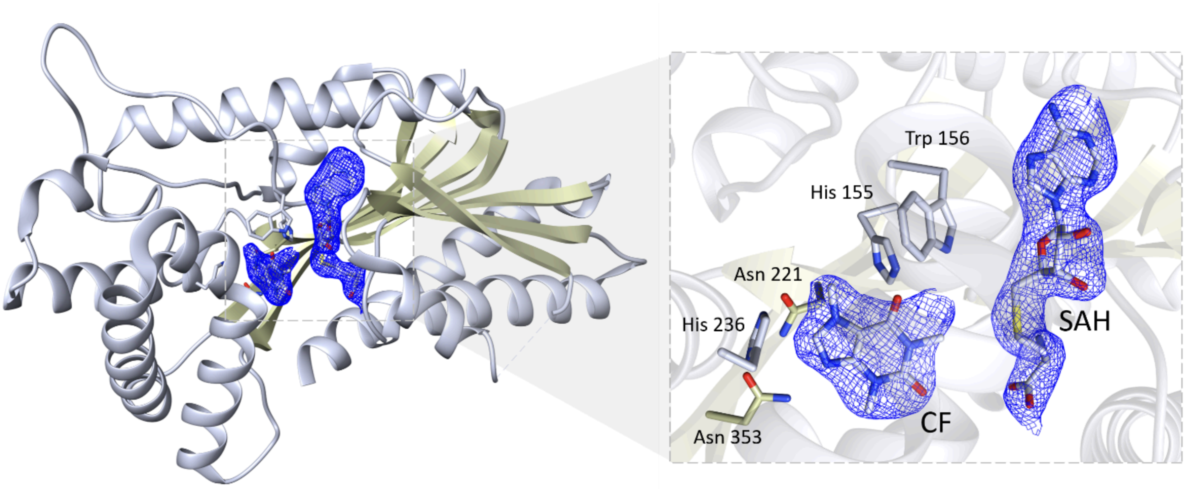

**Supplementary Figure 7. Crystal structure of IpCS3 displaying a difference Fourier map (Fo–Fc) contoured to 2.0 σ (blue) showing bound SAH and CF**. Relevant residues in IpCS3 for ligand recognition are displayed as lines with carbon atoms colored in gray, while small molecules - caffeine (CF), theobromine (TB), and S-adenosyl-homocysteine (SAH) - are drawn as sticks and labeled. Color code for the rest of the atoms: nitrogen (blue), oxygen (red) and sulphur (yellow).

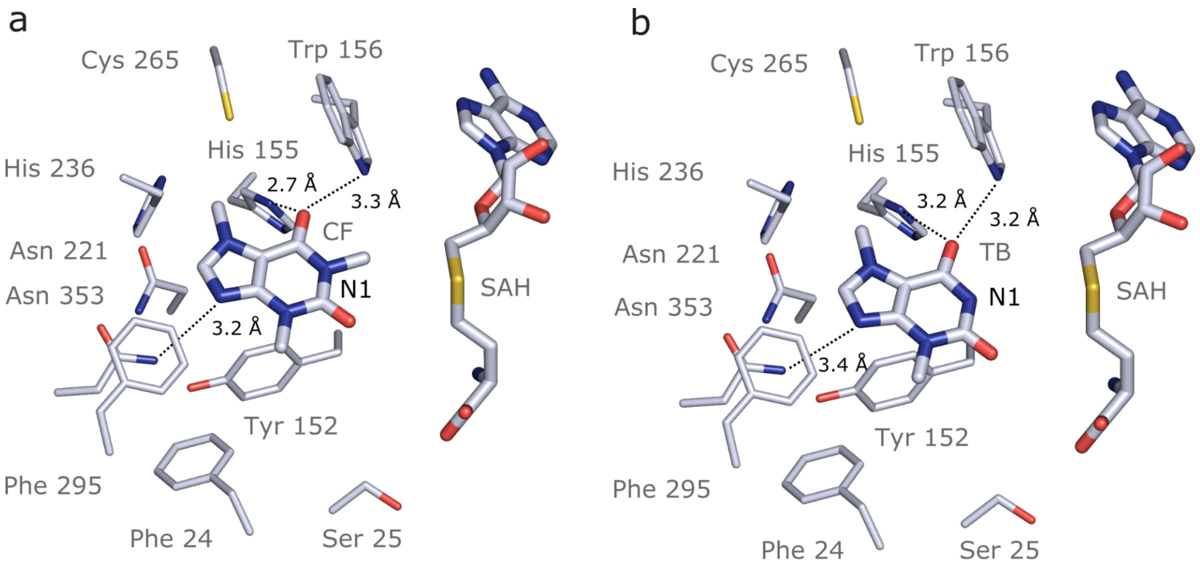

**Supplementary Figure 8. Theobromine and caffeine are oriented the same way in the active site of IpCS3. a,** IpCS3-CF complex (PDBID 8T2G). **b,** IpCS3-TB complex (docking model). Protein residues are displayed as lines with carbon atoms coloured in bluewhite while small molecules - theobromine (TB), caffeine (CF), and S-adenosyl-homocysteine (SAH) - are drawn as sticks. Colour code for the rest of the atoms: nitrogen (blue), oxygen (red) and sulphur (yellow). Hydrogen bond interactions are indicated as black dotted lines.

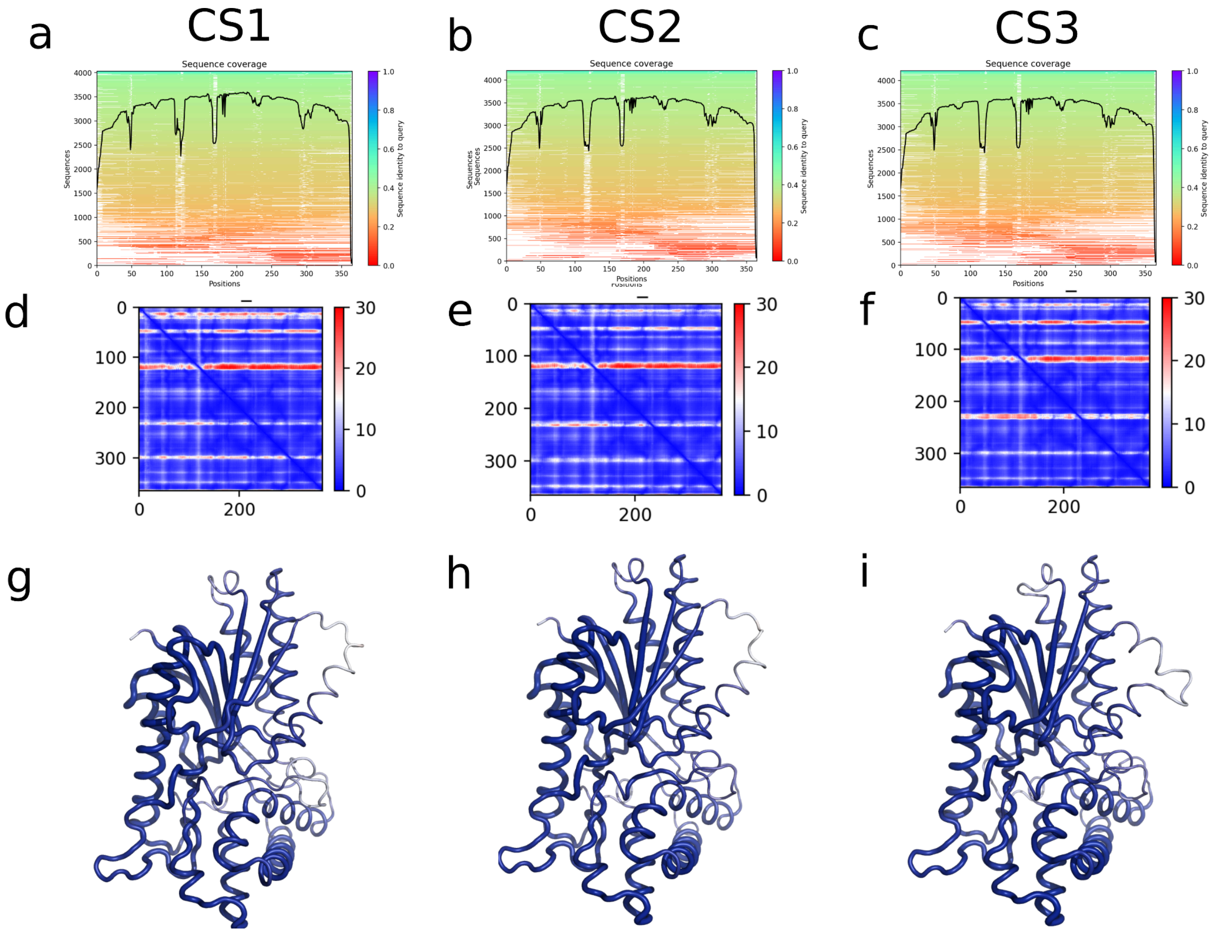

**Supplementary Figure 9. AlphaFold2-ColabFold Model Quality assessment of IpCS1, IpCS2 and IpCS3 models.** Sequence coverage of the MSA used for IpCS1 (a), IpCS2 (b) and IpCS3 (c). Alignment error for IpCS1 (d), IpCS2 (e) and IpCS3 (f). pLDDT score of IpCS1 (g), IpCS2 (h) and IpCS3 (i).

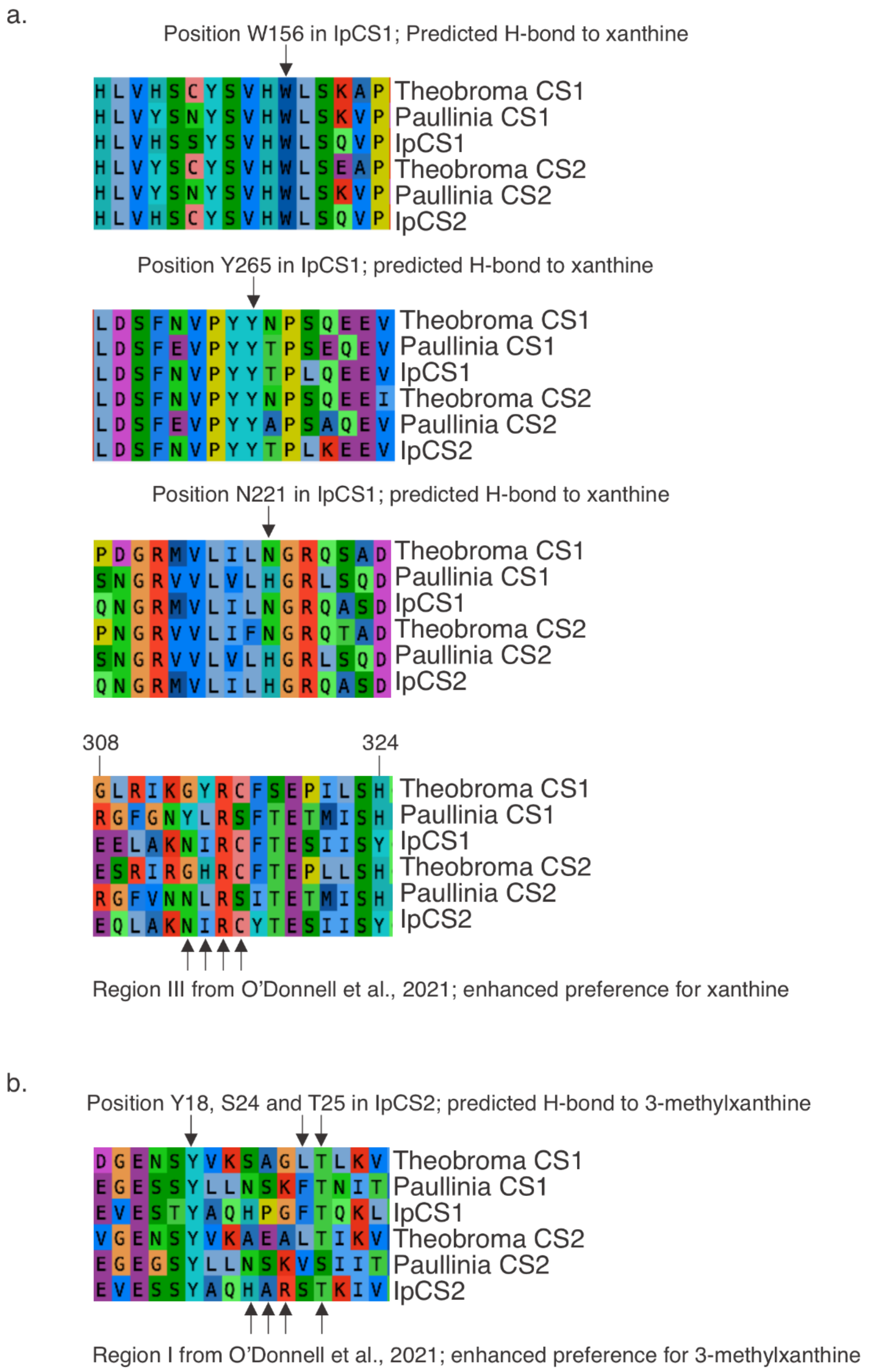

**Supplementary Figure 10. Comparative alignments of CS1 and CS2 show convergent changes predicted to participate in substrate binding and promote methylation preference switches. a,** comparison of homologous regions involved with xanthine binding. **b,** comparison of homologous regions involved with 3-methylxanthine binding.

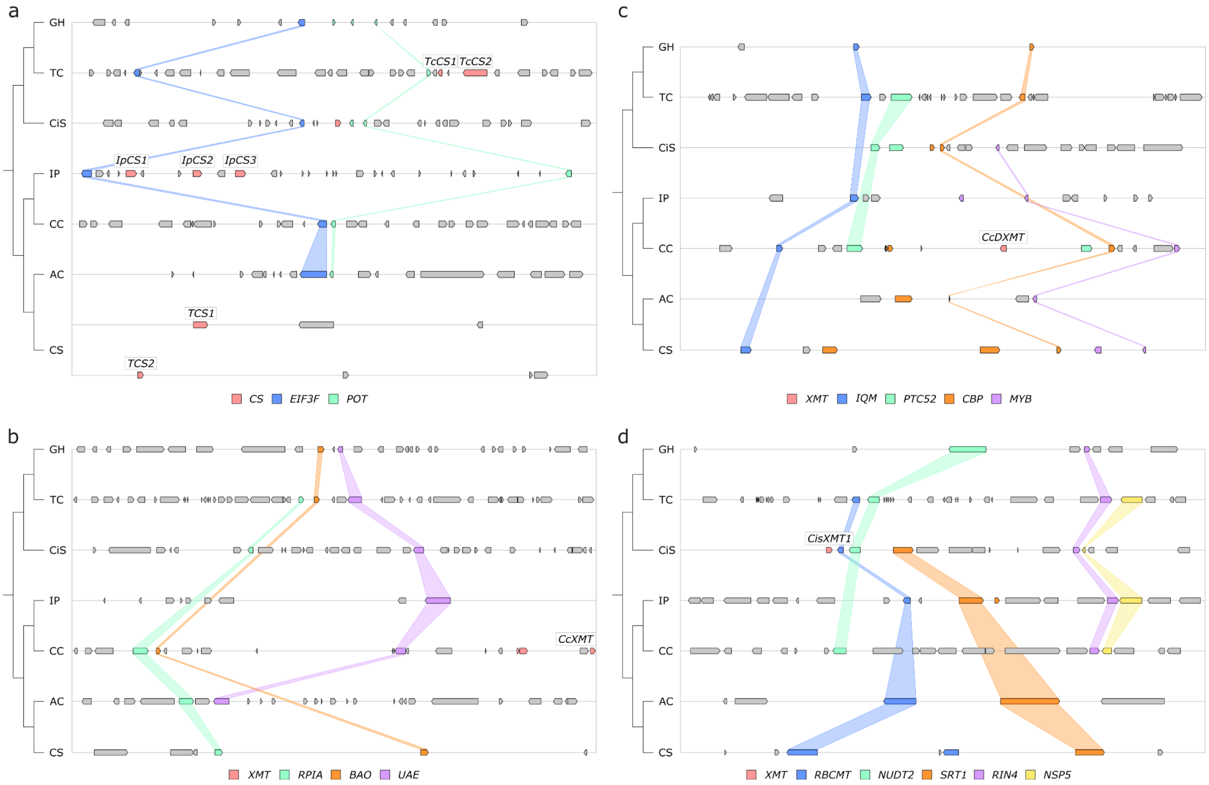

**Supplementary Figure 11. Only CS genes are available for co-option and utilisation for xanthine alkaloid biosynthesis in yerba mate. a,** Synteny-based analysis of the CS genomic region for seven angiosperm taxa. **b, c, d,** Synteny-based analyses of the XMT genomic regions for seven angiosperm taxa. Angiosperm taxa: GH, *Gossypium hirsutum*; TC, *Theobroma cacao*; CiS, *Citrus sinensis*; IP, *Ilex paraguariensis*; CC, *Coffea canephora*; AC, *Actinidia chinensis*; CS, *Camellia sinensis*. Genes: CS, caffeine synthase-type enzyme; EIF3F, eukaryotic translation initiation factor 3 subunit F; POT, proton-dependent oligopeptide transporter; XMT, xanthine methyltransferase-type enzyme; RPIA, ribose 5-phosphate isomerase A; BAO, beta-amyrin 28-oxidase-like; UAE, UDP-arabinose 4-epimerase 1-like; IQM, IQ domain-containing protein; PTC52, protochlorophyllide-dependent translocon component 52; CBP, calcium binding protein; MYB, MYB transcription factor; RBCMT, ribulose-1,5 bisphosphate carboxylase/oxygenase large subunit N-methyltransferase; NUDT2, nudix hydrolase 2-like, SRT1, NAD-dependent protein deacetylase; RIN4, RPM1 interacting protein 4; NSP5, nitrile specifier protein 5.

**Supplementary Tables**

Supplementary Table 1. Statistics of the genome sequencing data of yerba mate.

| Library | Number of reads | Read length | Total length | Coverage |
|---|---|---|---|---|
| Pair-end 350 bp #1 | 360,653,408 | 101 | 36.4 Gbp | 21.8 X |
| Pair-end 350 bp #2 | 368,746,464 | 101 | 37.2 Gbp | 22.3 X |
| Pair-end 550 bp | 356,261,246 | 101 | 36 Gbp | 21.5 X |
| Mate-pair 3 Kbp #1 | 415,398,586 | 101 | 30.3 Gbp | 18.2 X |
| Mate-pair 3 Kbp #2 | 410,588,934 | 101 | 30 Gbp | 17.9 X |
| Mate-pair 3 Kbp #3 | 343,059,350 | 101 | 25 Gbp | 15 X |
| Mate-pair 8 Kbp | 393,202,256 | 101 | 34.6 Gbp | 20.7 X |
| Mate-pair 12 Kbp | 415,478,776 | 101 | 33.7 Gbp | 20.1 X |
| PacBio long-reads | 19,514,627 | 50 bp - 61 Kbp | 77.5 Gbp | 49.3 X |
| Total | | | 341 Gbp | 207.8 X |

Supplementary Table 2. Statistics of the genome assembly of yerba mate.

| Metric | Value |
|---|---|
| # scaffolds (>= 1000 bp) | 10,611 |
| # scaffolds (>= 5000 bp) | 9,343 |
| # scaffolds (>= 10000 bp) | 8,951 |
| # scaffolds (>= 25000 bp) | 5,944 |
| # scaffolds (>= 50000 bp) | 2,595 |
| Total length (>= 50000 bp) | 887,124,725 |
| # scaffolds | 10,611 |
| Largest scaffold | 7,402,063 |
| Total length | 1,064,802,823 |
| GC (%) | 36.33 |
| N50 | 510,878 |
| N75 | 132,523 |
| L50 | 506 |
| L75 | 1,461 |
| # N's per 100 kbp | 1,976.99 |

Supplementary Table 3. Classification and distribution of repetitive DNA elements in yerba mate.

| | Number | Length occupied (bp) | Percentage of the genome (%) |
|---|---|---|---|
| Class I retrotransposons | 421,599 | 385,714,532 | 36.22 % |
| SINEs: | 840 | 154,298 | 0.01 % |
| Penelope | 0 | 0 | 0.00 % |
| LINEs: | 35,433 | 17,109,207 | 1.61 % |
| CRE/SLACS | 0 | 0 | 0.00 % |
| L2/CR1/Rex | 575 | 135,549 | 0.01 % |
| R1/LOA/Jockey | 443 | 76,937 | 0.01 % |
| R2/R4/NeSL | 0 | 0 | 0.00 % |
| RTE/Bov-B | 8,599 | 2,126,765 | 0.20 % |
| L1/CIN4 | 25,816 | 14,769,956 | 1.39 % |
| LTR retrotransposons: | 385,326 | 368,451,027 | 34.60 % |
| BEL/Pao | 709 | 266,632 | 0.03 % |
| Ty1/Copia | 98,237 | 67,631,136 | 6.35 % |
| Gypsy/DIRS1 | 216,472 | 274,526,515 | 25.78 % |
| Retroviral | 0 | 0 | 0.00 % |
| Class II DNA transposons | 45,427 | 19,116,209 | 1.80 % |
| hobo-Activator | 21,335 | 6,378,850 | 0.60 % |
| Tc1-IS630-Pogo | 0 | 0 | 0.00 % |
| En-Spm | 0 | 0 | 0.00 % |

Supplementary Table 3. (Continued)

| | | | |
|---|---|---|---|
| MuDR-IS905 | 0 | 0 | 0.00 % |
| PiggyBac | 0 | 0 | 0.00 % |
| Tourist/Harbinger | 5,870 | 2,846,548 | 0.27 % |
| Others | 0 | 0 | 0.00 % |
| Unclassified: | 990,080 | 269,430,122 | 25.30% |
| Total interspersed repeats: | 674,260 | 863 | 63.32 % |
| Small RNA: | 4,362 | 718,762 | 0.07 % |
| Satellites: | 0 | 0 | 0.00 % |
| Simple repeats: | 185,507 | 7,911,080 | 0.74 % |
| Low complexity: | 31,856 | 1,606,255 | 0.15 % |

Supplementary Table 4. Detail of yerba mate tRNA and anti-codon nucleotide sequences.

| tRNA genes | Anti-codon counts | | | | | | Total No. of tRNAs |
|---|---|---|---|---|---|---|---|
| POLAR | | | | | | | |
| Asparagine (Asn) | GTT (36) | ATT (0) | | | | | 36 |
| Cysteine (Cys) | GCA (22) | ACA (0) | | | | | 22 |
| Glutamine (Gln) | TTG (13) | CTG (10) | | | | | 23 |
| Glycine (Gly) | GCC (32) | TCC (11) | CCC (8) | ACC (0) | | | 51 |
| Serine (Ser) | GCT (20) | TGA (20) | AGA (15) | CGA (5) | GGA (5) | ACT (0) | 65 |
| Threonine (Thr) | TGT (11) | AGT (16) | GGT (6) | CGT (2) | | | 35 |
| Tyrosine (Tyr) | GTA (17) | ATA (0) | | | | | 17 |
| NON-POLAR | | | | | | | |
| Alanine (Ala) | AGC (12) | CGC (4) | TGC (11) | GGC (0) | | | 27 |
| Isoleucine (Ile) | AAT (14) | TAT (6) | GAT (2) | | | | 22 |
| Leucine (Leu) | CAA (23) | AAG (10) | CAG (4) | TAG (8) | TAA (6) | GAG (0) | 51 |
| Methionine (Met) | CAT (55) | | | | | | 55 |
| Phenylalanine (Phe) | GAA (30) | AAA (2) | | | | | 32 |
| Proline (Pro) | AGG (10) | TGG (28) | CGG (4) | GGG (0) | | | 42 |
| Tryptophan (Trp) | CCA (31) | | | | | | 31 |
| Valine (Val) | AAC (11) | GAC (10) | CAC (9) | TAC (7) | | | 37 |
| POSITIVELY CHARGED | | | | | | | |
| Arginine (Arg) | ACG (15) | TCT (14) | CCT (7) | CCG (6) | TCG (6) | GCG (3) | 51 |
| Histidine (His) | GTG (25) | ATG (2) | | | | | 27 |
| Lysine (Lys) | CTT (10) | TTT (17) | | | | | 27 |
| NEGATIVELY CHARGED | | | | | | | |
| Aspartic acid (Asp) | GTC (39) | ATC (1) | | | | | 40 |
| Glutamic acid (Glu) | CTC (14) | TTC (21) | | | | | 35 |

Supplementary Table 4. (Continued)

| | | | | |
|---|---|---|---|---|
| Selenocysteine tRNAs | TCA (0) | | | 0 |
| Possible suppressor tRNAs | CTA (0) | TTA (1) | TCA (1) | 2 |
| tRNAs with undetermined isotypes | | | | 11 |
| Predicted pseudogenes | | | | 76 |

Supplementary Table 5. miRNA families predicted in the yerba mate genome.

| miRNA | Functional involvement in other eudicot plants |
|---|---|
| miR156 | Seed growth and development[12,13] |
| | Fruit development[14] |
| | Drought/cold stress[15,16] |
| miR159 | Growth and development[17] |
| | Phase change from vegetative to reproductive growth[18] |
| | Lipid and protein accumulation[19] |
| | Drought stress[20] |
| miR160 | Growth and development[21,22] |
| | Fibrous root and storage root development[23] |
| | Drought stress[24] |
| miR162_2 | Storage root initiation and development[23] |
| miR164 | Lateral root and leaf development[25] |
| | Fibrous root and storage root development[23] |
| | Seed development[12] |
| | Drought stress[26] |
| miR166 | Seed development[12] |
| | Fibrous root and storage root development[23] |
| | Drought stress[20] |
| | Disease resistance[27] |
| miR167_1 | Growth and development[17] |
| | Drought/cold stress[20,28] |
| miR168 | Development[22] |
| | Resistance to fire blight[29] |
| miR169_2; miR169_5 | Drought/cold/salt stress[30–33] |
| miR171_1; miR171_2 | Development[34,35] |
| | Lipid and protein accumulation[19] |
| miR172 | Development[36] |
| | Starch biosynthesis[37] |
| | Drought/cold stress[32] |
| miR390 | Drought stress[31] |
| | Leaf morphology[38] |
| miR394 | Drought/salt stress[39] |

Supplementary Table 5. (Continued)

| | |
|---|---|
| miR395 | Low sulfate response[40] |
| miR396 | Seed development[41] |
| | Starch biosynthesis[37] |
| | Drought/salt stress[31,42] |
| miR397 | Drought/cold stress[32] |
| miR398 | Fibrous root and storage root development[23] |
| | Salt stress[33] |
| miR399 | Phosphate homeostasis[40,43] |
| | Shoot to root transport[43] |
| miR403 | Drought stress[31] |
| miR405 | Transposon derived[44] |
| miR408 | Tolerance to Boron deficiency[45] |
| | Cold stress[46] |
| | Response to wounding and topping[47] |
| miR473 | Metabolism[48] |
| | Stress response[49] |
| miR474 | Drought stress[50] |
| miR475 | Metabolism[48] |
| miR477 | Starch biosynthesis[51] |
| miR530 | Disease resistance[52] |
| miR1023 | Disease resistance[53] |
| miR1446 | Stress response[54] |

Supplementary Table 6. miRNA targets predicted in the yerba mate genome.

| Targets IDs | Description | miR159 | miR164 | miR167_1 | miR168 | miR169_2 | miR169_5 | miR171_1 | miR171_2 | miR390 | miR394 | miR396 | miR397 | miR398 | miR403 |
|---|---|---|---|---|---|---|---|---|---|---|---|---|---|---|---|
| ILEXPARA_008283 | Uncharacterized protein | ✷ | | | | | | | | | | | | | |
| ILEXPARA_029002 | Uncharacterized protein | ✷ | | | | | | | | | | | | | |
| ILEXPARA_031381 | Uncharacterized protein | ✷ | | | | | | | | | | | | | |
| ILEXPARA_043376 | Uncharacterized protein | ✷ | | | | | | | | | | | | | |
| ILEXPARA_013180 | *ileS*, isoleucine tRNA ligase | ✷ | | | | | | | | | | | | | |
| ILEXPARA_000910 | myb-like transcription factor | ✷ | ✷ | | | | | | | | | | | | |
| ILEXPARA_028644 | Uncharacterized protein | | ✷ | | | | | | | | | | | | |
| ILEXPARA_005969 | Putative membrane protein | | ✷ | | | | | | | | | | | | |
| ILEXPARA_048009 | *panC*, pantothenate (vitamin B5) synthetase | | ✷ | | | | | | | | | | | | |
| ILEXPARA_018064 | *arf*, auxin response factor | | | ✷ | | | | | | | | | | | |
| ILEXPARA_019275 | Uncharacterized protein | | | ✷ | | | | | | | | | | | |
| ILEXPARA_024153 | Uncharacterized protein | | | ✷ | | | | | | | | | | | |
| ILEXPARA_035190 | Uncharacterized protein | | | ✷ | | | | | | | | | | | |
| ILEXPARA_016483 | Hypothetical protein | | | | ✷ | | | | | | | | | | |
| ILEXPARA_029421 | *GCP4*, gamma tubulin complex protein 4 | | | | | ✷ | ✷ | | | | | | | | |
| ILEXPARA_047849 | *GOLS1*, galactinol synthase 1 | | | | | ✷ | ✷ | | | | | | | | |
| ILEXPARA_003987 | *NACK1*, kinesin-like protein | | | | | ✷ | ✷ | | | | | | | | |
| ILEXPARA_044341 | Uncharacterized protein | | | | | ✷ | | | | | | | | | |

Supplementary Table 6. (Continued)

| | | | | | | | | | | | | | | |
|---|---|---|---|---|---|---|---|---|---|---|---|---|---|---|
| ILEXPARA_005359 | Uncharacterized protein | | | | | ✷ | | | | | | | | |
| ILEXPARA_035716 | *RABE1C*, ras-related protein | | | | | ✷ | | | | | | | | |
| ILEXPARA_010316 | *MAPK*, mitogen activated protein kinase | | | | | ✷ | ✷ | | | | | | | |
| ILEXPARA_032923 | Uncharacterized protein | | | | | ✷ | ✷ | | | | | | | |
| ILEXPARA_008149 | Uncharacterized protein | | | | | ✷ | ✷ | | | | | | | |
| ILEXPARA_048631 | Protein kinase | | | | | ✷ | ✷ | | | | | | | |
| ILEXPARA_008152 | Uncharacterized protein | | | | | | | ✷ | | | | | | |
| ILEXPARA_023090 | *NAGK*, N-acetyl-D-glucosamine kinase | | | | | | | ✷ | | | | | | |
| ILEXPARA_024088 | RNA-binding (RRM/RBD/RNP motif) family protein | | | | | | | ✷ | | | | | | |
| ILEXPARA_023716 | Endoglucanase | | | | | | | ✷ | | | | | | |
| ILEXPARA_042182 | Uncharacterized protein | | | | | | | ✷ | | | | | | |
| ILEXPARA_021515 | Pentatricopeptide repeat (PPR) protein | | | | | | | | ✷ | | | | | |
| ILEXPARA_004925 | Uncharacterized protein | | | | | | | | | ✷ | | | | |
| ILEXPARA_045111 | Rotamase *FKBP 1* | | | | | | | | | ✷ | | | | |
| ILEXPARA_013832 | *ABCC2*, ABC transporter C family member 2 protein | | | | | | | | | | ✷ | | | |
| ILEXPARA_039828 | *guaA*, GMP synthase | | | | | | | | | | ✷ | | | |
| ILEXPARA_028274 | Hypothetical protein | | | | | | | | | | ✷ | | | |
| ILEXPARA_024538 | *RPT6A*, regulatory particle triple-A ATPase 6A | | | | | | | | | | | ✷ | | |
| ILEXPARA_031387 | Uncharacterized protein | | | | | | | | | | | | | ✷ |

Supplementary Table 6. (Continued)

| | | | | | | | | | | | | | | | |
|---|---|---|---|---|---|---|---|---|---|---|---|---|---|---|---|
| ILEXPARA_043757 | Uncharacterized protein | | | | | | | | | | | | | ✷ | |
| ILEXPARA_005297 | Uncharacterized protein | | | | | | | | | | | | | ✷ | |
| ILEXPARA_012032 | Uncharacterized protein | | | | | | | | | | | | | ✷ | |
| ILEXPARA_9682 | *OST1B*, oligosaccharyltransferase 1B | | | | | | | | | | | | | | ✷ |

Supplementary Table 7. Apparent enzyme kinetic parameter estimates for yerba mate caffeine biosynthetic enzymes with selected substrates.

| Enzyme (substrate) | $K_M$ (µM) | $k_{cat}$ (1/sec) | $k_{cat}/K_M$ ($s^{-1}M^{-1}$) |
|---|---|---|---|
| IpCS1 (X) | 85.05 | 0.0009 | 10.11 |
| IpCS2 (3X) | 197.08 | 0.0031 | 15.77 |
| IpCS3 (TB) | 151.19 | 0.0029 | 19.36 |

Supplementary Table 8. Data collection and refinement statistics of IpCS3 structure bound to SAH and caffeine.

| IpCS3 in complex with SAH and caffeine | |
|---|---|
| **PDB** | 8UZD |
| **Data collection** | |
| Wavelength (Å) | 0.9786 |
| Resolution (Å) | 2.72 |
| Resolution Range[a] | 37.00- 2.72 |
| | (2.82 - 2.72) |
| Space group | P $4_1$ $2_1$ 2 |
| Cell dimensions | |
| *a*, *b*, *c* (Å) | 82.67, 82.67, 226.09 |
| α, β, γ (°) | 90.00, 90.00, 90.00 |
| Total reflections | 43,818 |
| Unique reflections | 21,910 |
| Multiplicity[a] | 2.0 (2.0) |
| Completeness (%)[a] | 99.89 (100.00) |
| $\langle I / \sigma I \rangle$[a] | 25.79 (2.87) |
| $R_{merge}$[a,b] (%) | 0.0223 (0.2168) |
| $R_{meas}$ (%)[a] | 0.0315 (0.3066) |
| $CC_{1/2}$[a] | 0.999 (0.878) |
| **Refinement** | |
| Resolution (Å) | 2.72 |
| No. reflections | 21,909 |
| $R_{work}$[c] / $R_{free}$[d] | 0.194/0.248 |
| No. atoms | |

Supplementary Table 8. (Continued)

| | |
|---|---|
| Protein | 5,216 |
| CFF + SAH | 80 |
| Water | 48 |
| *B*-factors | |
| Protein | 63.38 |
| CFF + SAH | 84.48 |
| Water | 48.19 |
| Bond lengths (Å) | 0.004 |
| Bond angles (°) | 1.112 |

[a] Numbers in parentheses refer to the highest resolution shell.
[b] $R_{merge} = \Sigma |I_i - <I_i>| / \Sigma I_i$ where $I_i$ = the intensity of the $i$th reflection and $<I_i>$ = mean intensity.
[c] $R_{work} = \Sigma |F_o - F_c| / \Sigma |F_o|$, where $F_o$ and $F_c$ are the observed and calculated structure factors, respectively.
[d] $R_{free}$ was calculated as for $R_{work}$, but on a test set comprising 5% of the data excluded from refinement.

Supplementary Table 9. Accession numbers of SABATH sequences used for phylogenetic analysis in Figure 3.

| Accession Number | Gene | Species |
|---|---|---|
| ABV91100.1 | CCMT1 | *Ocimum basilicum* |
| NP_001406848.1 | IAMT1 | *Oryza sativa subsp. japonica* |
| NP_200336.1 | IAMT1 | *Arabidopsis thaliana* |
| KAH9652080.1 | IAMT1 | *Citrus sinensis* |
| XP_002298843.1 | IAMT1 | *Populus trichocarpa* |
| NP_194372.2 | GAMT1 | *Arabidopsis thaliana* |
| NP_200441.2 | GAMT2 | *Arabidopsis thaliana* |
| NP_190072.1 | FAMT | *Arabidopsis thaliana* |
| gnl\|onekp\|DAAD_scaffold_2041891 | FAMT | *Ardisia revoluta* |
| B2KPR3.1 | LAMT | *Catharanthus roseus* |
| XP_015624979.1 | BSMT | *Oryza sativa subsp. japonica* |
| D9J0Z7.1 | AAMT1 | *Zea mays* |
| NP_187755.2 | BSMT1 | *Arabidopsis thaliana* |
| AAP57211.1 | BSMT1 | *Arabidopsis lyrata subsp. lyrata* |
| gnl\|onekp\|VKGP_scaffold_2120585 | CS | *Geranium carolinianum* |
| gnl\|onekp\|YGCX_scaffold_2142952 | CS | *Geranium maculatum* |
| EC774687 | CS0 | *Paullinia cupana var. sorbilis* |
| EC778019 | CS2 | *Paullinia cupana var. sorbilis* |
| EC766748 | CS1 | *Paullinia cupana var. sorbilis* |
| DAA64605.1 | CS | *Paullinia cupana var. sorbilis* |

Supplementary Table 9. (Continued)

| | | |
|---|---|---|
| gnl\|onekp\|VFFP_scaffold_2043613 | CS | *Acer negundo* |
| gnl\|onekp\|WAXR_scaffold_2040211 | CS | *Litchi chinensis* |
| KDO69071.1 | CS | *Citrus sinensis* |
| GBCV01000539.1 | CS | *Mangifera indica* |
| gnl\|onekp\|YUOM_scaffold_2035237 | CS | *Rhus radicans* |
| gnl\|onekp\|BCAA_scaffold_2069913 | CS | *Kirkia wilmsii* |
| gnl\|onekp\|FCCA_scaffold_2008319 | CS | *Boswellia sacra* |
| KAJ4720014.1 | CS | *Melia azedarach* |
| gnl\|onekp\|WMUK_scaffold_2093724 | CS | *Schizolaena* sp. |
| gnl\|onekp\|ATFX_scaffold_2034890 | CS | *Muntingia calabura* |
| BAE79730.1 | BTS1 | *Theobroma cacao* |
| A0A061FKL9.1 | CS1 | *Theobroma cacao* |
| A0A061FKM4.1 | CS2 | *Theobroma cacao* |
| Q68CM3.1 | CS2 | *Camellia sinensis* |
| Q9FZN8.1 | CS1 | *Camellia sinensis* |
| gnl\|onekp\|ADHK_scaffold_2052625 | CS | *Galax urceolata* |
| gnl\|onekp\|PPPZ_scaffold_2019622 | CS | *Rhododendron scopulorum* |
| gnl\|onekp\|ODDO_scaffold_2102454 | CS | *Ardisia humilis* |
| gnl\|onekp\|WMUK_scaffold_2020889 | CS | *Schizolaena* sp. |
| Q9SPV4.1 | SAMT | *Clarkia breweri* |

Supplementary Table 9. (Continued)

| | | |
|---|---|---|
| AAN40745.1 | SAMT | *Antirrhinum majus* |
| CAI05934.1 | SAMT | *Hoya carnosa* |
| ACZ55216.1 | SAMT | *Nicotiana suaveolens* |
| NP_001289539.1 | SAMT | *Nicotiana sylvestris* |
| ACZ55219.1 | SAMT | *Nicotiana alata* |
| EF472972.1 | SAMT | *Datura wrightii* |
| BAB39396.1 | SAMT | *Atropa belladonna* |
| KAH9711297.1 | SAMT | *Citrus sinensis* |
| ACZ55224.1 | NAMT | *Nicotiana gossei* |
| AJ628349.1 | BSMT1 | *Nicotiana suaveolens* |
| NA | NAMT | *Nicotiana suaveolens* |
| ACZ55223.1 | BSMT2 | *Nicotiana sylvestris* |
| ACZ55220.1 | BSMT2 | *Nicotiana alata* |
| ACZ55217.1 | BSMT2 | *Nicotiana suaveolens* |
| AAG23343.1 | JMT | *Arabidopsis thaliana* |
| XP_002307671.1 | JMT | *Populus trichocarpa* |
| XP_004291853.1 | JMT2 | *Fragaria vesca subsp. vesca* |
| XP_004291852.1 | JMT1 | *Fragaria vesca subsp. vesca* |
| Q9FYZ9.1 | BAMT | *Antirrhinum majus* |
| KDO50937.1 | XMT1 | *Citrus sinensis* |

Supplementary Table 9. (Continued)

| | | |
|---|---|---|
| KDO40396.1 | XMT2 | *Citrus sinensis* |
| XM_006469387.4 | XMTB | *Citrus sinensis* |
| XM_024190084.1 | XMTA | *Citrus sinensis* |
| AFV60438.1 | DXMT1 | *Coffea arabica* |
| XP_027086771.1 | XMT1 | *Coffea arabica* |
| NP_001392358.1 | MXMT1 | *Coffea arabica* |
| Q84PP7.1 | MXMT2 | *Coffea arabica* |
